# Dynamic Tracking of Native Precursors in Adult Mice

**DOI:** 10.1101/2024.04.02.587737

**Authors:** Suying Liu, Sarah E. Adams, Haotian Zheng, Juliana Ehnot, Seul K. Jung, Greer Jeffrey, Theresa Menna, Louise E. Purton, Hongzhe Lee, Peter Kurre

**Author notes:** Correspondence: Correspondence to Peter Kurre.

## Abstract

Hematopoietic dysfunction has been associated with a reduction in the number of active precursors. However, precursor quantification at homeostasis and under diseased conditions is constrained by the scarcity of available methods. To address this issue, we optimized a method for quantifying a wide range of hematopoietic precursors. Assuming the random induction of a stable label in precursors following a binomial distribution, estimates depend on the inverse correlation between precursor numbers and the variance of precursor labeling among independent samples. Experimentally validated to cover the full dynamic range of hematopoietic precursors in mice (1 to 10^5^), we utilized this approach to demonstrate that thousands of precursors, which emerge after modest expansion during fetal-to-adult transition, contribute to native and perturbed hematopoiesis. We further estimated the number of precursors in a mouse model of Fanconi Anemia, showcasing how repopulation deficits can be classified as autologous (cell proliferation) and non-autologous (lack of precursor). Our results support an accessible and reliable approach for precursor quantification, emphasizing the contemporary perspective that native hematopoiesis is highly polyclonal.

## Introduction

Continuous self-renewal and differentiation of hematopoietic stem and progenitor cells (HSPCs) is fundamental to blood production. Even though rare cases exist where a single HSPC clone supports hematopoiesis, the HSPC population contributing to homeostatic hematopoiesis is usually highly polyclonal [1,2]. For example, the number of HSPCs actively participating in white blood cell production was estimated to range from 20,000 to 200,000 in humans [3,4]. Conversely, a decline in the number of active hematopoietic precursors has been linked to hematopoietic dysfunction. For example, in humans older than 70 years of age, hematopoietic dysfunction coincides with an abrupt reduction in the HSPCs population actively contributing to blood production [4].

These observations underscore the association between a low number of active hematopoietic precursors and hematological function. Yet, few methods are suitable for quantifying active hematopoietic precursors in a native environment [3–8]. In mice, methods utilizing *in situ* barcodes often suffer from barcode homoplasy, preventing total precursor number estimation [9,10]. In humans, estimations based on somatic mutations and computational modeling carry a high degree of uncertainty [3,4]. The absence of precise precursor quantification methods in their natural environment hampers the study of precursor numbers across different conditions and their potential use as predictive functional markers.

In mice, the quantification of developmental hematopoietic precursors has been achieved using animals engineered with a Confetti cassette [11]. This cassette can randomly recombine and express one of four fluorescence proteins (FPs, being RFP, CFP, YFP, or GFP) upon Cre induction, resulting in variability in Confetti expression pattern in mice that inversely correlates with precursor numbers (Figure 1A) [11]. While suitable for developmental hematopoiesis, this method has limited linear range (50-2500), impeding its potential application to adult hematopoiesis or clonally restricted hematopoiesis. To measure hematopoietic precursors in various conditions, expansion of the detection range is desired.

**Figure 1.**
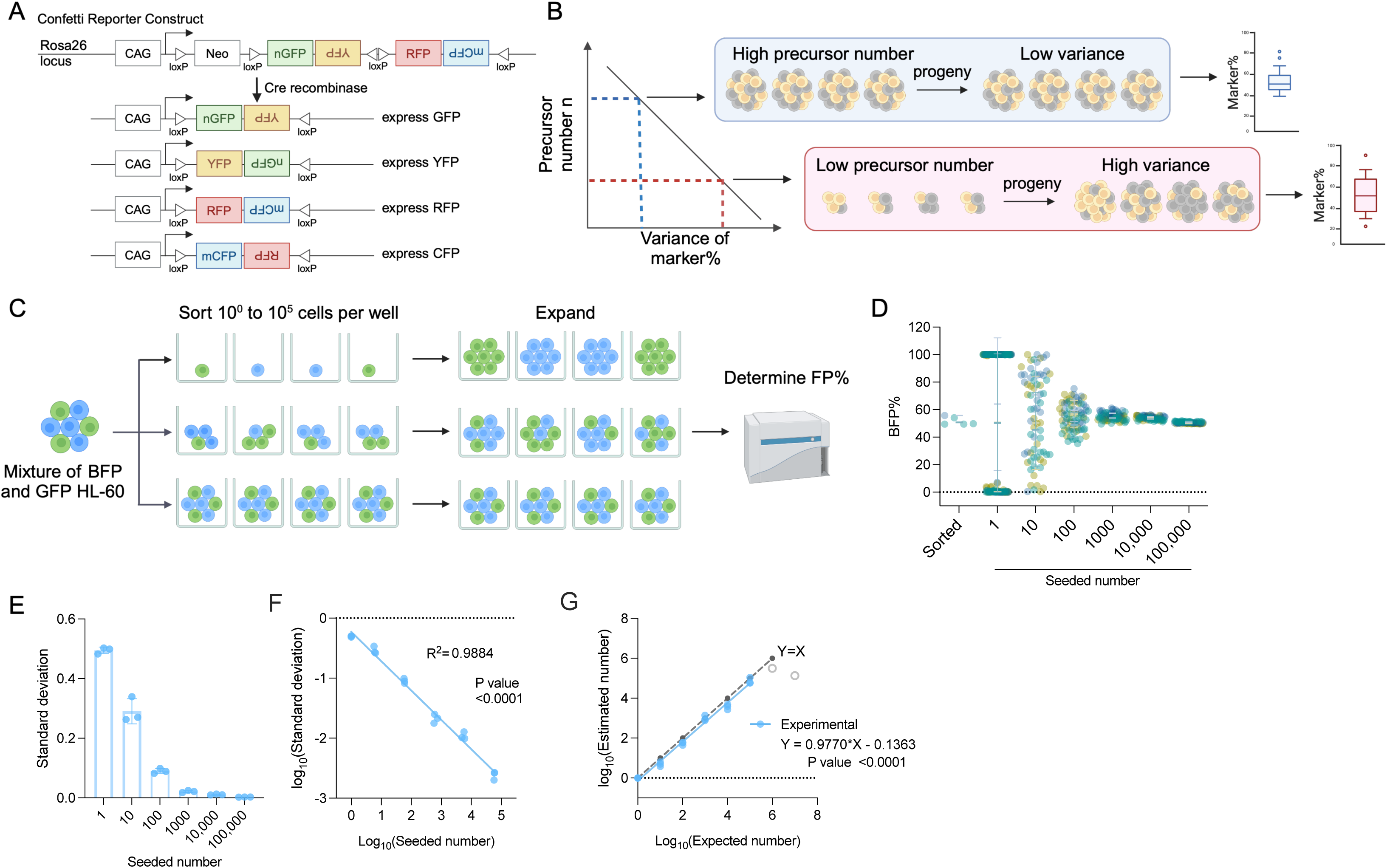
The Principle of Inverse Correlation between Variance of FP% and precursor numbers. (A) Schematic of the Confetti cassette at the mouse *Rosa26* locus. Sequences of four fluorescence protein are interspaced by loxP sites. Upon Cre-expression, the Confetti cassette will recombine and express one of the four fluorescence proteins. (B) The inverse relationship between variance of markers and precursor numbers. The distribution of FP% is determined in the progeny to estimate the number of precursors. (C) Workflow to validate the correlation formula between variance of FP% and precursor numbers by a two-color cell system. (D) BFP frequencies in wells seeded with 1 to 100,000 HL-60. Each dot represents a well. Two replicates were shown. Each replicate consists of at least 20 wells per seeded number. Error bars represent mean ± SD. (E) The standard deviation of BFP% in wells seeded with various numbers of HL-60. Error bars represent mean ± SD. N = 3 replicates. (F) The correlation between seeded numbers and standard deviation of BFP%. N = 3 replicates. (G) The correlation between seeded numbers and the numbers estimated with standard deviation of BFP% and equation 2. N = 3 replicates except for seeded numbers 10^6^ and 10^7^ (grey circles). For 10^6^ and 10^7^, N= 1 replicates. For (A-C), images were created with Biorender.

Interestingly, the random induction of Confetti colors in HSPCs bears similarity with X-chromosome inactivation (XCI). In XCI, one of the X chromosomes is randomly inactivated in precursor cells, a process that adheres to a binomial distribution. Formula derivation from binomial distribution implies that a higher XCI variance among individuals correlates with a smaller number of precursors at the time of inactivation (Figure 1B). As XCI is faithfully maintained in the progeny, this correlation has been used to estimate precursor numbers from mature blood cells in human and mouse, leading to the discovery of the first mutation (Tet2) that contributes to clonal hematopoiesis [7,12–14]. Nonetheless, XCI occurs exclusively in females, limiting its applicability in males. Additionally, XCI takes place during early development when few precursors are present, resulting in low sensitivity in adulthood [15].

Building on insights from Confetti mice and XCI studies, we investigated whether random induction of Confetti FPs in precursor cells can be similarly modeled by a binomial distribution, whereby the variance of FPs inversely correlates with precursor numbers. Based on this relationship, we asked if we could broaden the measurable range of precursor number, which would allow us to expand the existing correlation range.

Examining the premises of binomial distribution and setting some assumptions, we report that the random induction of FPs among a group of mice or cells can be modeled by a binomial distribution. Experimental validation establishes a broadened linear range, covering the full spectrum of precursor numbers (1∼10^5^) and overcoming the prior range limit. We leverage this correlation to probe the number of hematopoietic precursors at homeostasis, post-myeloablation, during developmental expansion, and in a mouse model of inherited bone marrow failure.

## Results

### Binomial distribution underlies the inverse linear correlation between FP% variance and the precursor numbers

To broaden the limited correlation range, we aimed to elucidate the mathematical relationship between variance and precursor numbers [11]. Inspired by XCI, we asked how the stochastic induction of Confetti FPs in precursors may satisfy the premises of binomial distribution [16,17]. We found that when we assume the number of precursors remain the same for the condition tested, then induction of FPs meets the required premises: (1) the number of precursors, denoted as *n*, remains constant within a group of mice or cells; (2) each observation (e.g. each precursor) is independent; (3) each observation yields one of two outcomes (e.g., it signifies the presence or absence of RFP in a precursor); (4) the probability of success, represented as *p* (e.g., the likelihood of a precursor expressing RFP upon Confetti induction), remains consistent across all observations.

Should the induction of a FP adhere to a binomial distribution, precursor numbers can be estimated using the following equation:

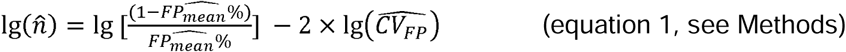

where 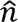 signifies the estimated number of precursors, 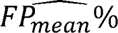 represents the estimated probability of a precursor being one given FP (e.g., 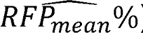), and 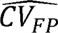 denotes the estimated coefficient of variance (CV) of one given FP (e.g., 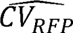). CV is the standard deviation divided by the mean, utilized by the empirical correlation formula.

The equation proposes a direct linear relationship between the precursor number and the CVs of individual FPs, supporting the empirical correlation formula [11]. Indeed, by analyzing a published dataset providing precursor numbers and the corresponding distribution of FPs, we found that the correlation between *n* and individual CVs of FP exhibits superior fit (>0.93), higher than the empirical correlation equation based on all three CVs (0.75) (Figure S1A) [11]. Moreover, according to equation 1, the y-intercept of correlation between precursor number and CV was influenced by FP_mean_%, while the slope was the same (-2) regardless of FP_mean_%. Given the unequal value of RFP_mean_%, CFP_mean_%, and YFP_mean_% in the published dataset, the correlation equations show significantly distinct y-intercepts (p<0.0001) (Figure S1A and B). In contrast, the slopes of these correlation equations were similar (p=0.49, with all 95% confidence intervals encompassing -2, the theoretical value) (Figure S1A).

In conclusion, the FP induction in precursor cells can be modeled by a binomial distribution with the assumption that precursor numbers are constant among a group of mice. This sets the mathematical basis for the inverse linear correlation between variance of FP% and the number of precursors.

### A broad range of precursor numbers correlates with variance of FP%

While the correlation formula derived from binomial distribution does not impose any range limitation, experimental errors may confound the measurement of variance. We next aimed to experimentally confirm this correlation. To streamline the calculation, we used standard deviation instead of CV to compute variance. The correlation between standard deviation and precursor numbers is expressed through the following equation:

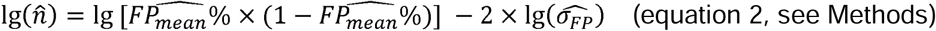

where 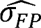 denotes the estimated standard deviation of a given FP% (e.g. 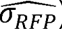).

Given this equation, 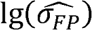 exhibits a linear and inverse correlation with 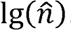. Indeed, 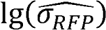, 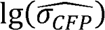 and 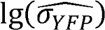 exhibited a linear and inverse correlation with 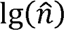 in the published data (Figure S1C) [11].

To simplify the validation, we used a two-color cell model HL-60 bearing one of the two FPs (BFP and GFP, wherein GFP represents non-BFP) (Figure 1C). Although the Confetti cassette recombination can generate one of four colors, our estimation is based on a single given FP (e.g. a precursor expresses RFP or not), making this simplification justifiable.

We first proved that neither BFP nor GFP HL-60 cells had a competitive growth advantage over the other, ensuring that HL-60 progeny mirrored the seeding population (Figure S1D). We next sorted one to 10^7^ cells into individual wells and allowed them to proliferate for at least three generations before assessing BFP% at the end of the culture (Figure 1C). We focus on cell range up to 10^5^ cells as 1.4×10^5^ non-HSC LSK and 5.2×10^3^ active HSCs were previously estimated in an adult female mouse, which corresponds to the highest possible precursor numbers in mice [18]. We considered the survival rate of cells in wells receiving a single HL-60 cell (53% to 60%), to ensure the accuracy of the number of cells seeded.

Consistent with the proposed correlation, as seeded numbers increase, BFP% standard deviation decreases (Figure 1D and E). The variance of BFP% reveals an inverse correlation with the seeding numbers (R^2^ = 0.996) (Figure 1F). After normalizing for cell survival rates, the calculated numbers closely align with the expected numbers from one to 10^6^ cells (Figure 1G). We had anticipated that measurement of small variance in high precursor numbers may be confounded by experimental errors. Indeed, at cell numbers larger than 10^6^, the standard deviation of BFP% and the expected number reached a plateau (Figure 1G, grey circles). To ensure accurate measurement without confounding errors, we suggest 10^5^ to be the upper limit for experimental measurement.

### Experimental practices for accurate precursor number measurement

It is crucial to ensure the accuracy of variance measured because variance of FP% is solely used for estimation. Based on the data from the two-color HL-60 model, we conclude that there are at least three experimental practices required for accurate cell estimates: (1) exclusion of outliers; (2) sufficient flow cytometry recorded events; (3) sufficient sample size per group. Both outliers and insufficient recorded events inaccurately inflate sample variance, leading to underestimation of precursor numbers (Figure S1E and F). We found that the minimum recorded events increase in tandem with the seeded number, in contrast to the reported 500-event threshold (Figure S1F) [11].

While having a small sample size per biological replicate is possible, the variability of estimates among replicates would be high (Figure S1G). Practically, achieving a large sample size (>20) is cumbersome in mice. To determine a feasible sample size for relatively accurate estimates, we resampled datapoints to calculate the variability reduction as sample size increases (Figure S1H). Our analyses reveal that five samples per group are sufficient to substantially minimize error, whereas an additional increase in sample size leads to marginal error reduction. We therefore use at least five mice per biological replicates for our precursor estimations.

### Inducible HSC-Scl-CreER^T^-targeted Confetti labeling in the active blood precursors at the adult stage

To label hematopoietic precursors and quantify their number *in vivo*, we considered the choice of Cre mouse line to be crossed with Confetti. Historically, hematopoietic precursors are thought to be hematopoietic stem cells (HSCs) capable of long-term repopulation. However, recent studies indicate that multipotent progenitors (MPPs) also contribute alongside HSCs in maintaining blood production [19–22]. Hence, both HSCs and MPPs should be labeled.

Among Cre mouse lines capable of labeling HSCs and MPPs simultaneously at the adult stage (such as Rosa26^CreERT2^, Mx1-Cre, and HSC-Scl-CreER^T^), we chose HSC-Scl-CreER^T^ because of its preference for labeling HSPCs (immunophenotypically defined as Lin^-^Sca-1^+^cKit^+^ (LSK)) [23]. To validate the specificity of HSC-Scl-CreER^T^, we generated mice possessing a single Confetti allele and homozygous HSC-Scl-CreER^T^ alleles. We then examine Confetti expression one day after a two-day tamoxifen administration. Consistent with prior reports of HSC-Scl-CreER^T^ activity, we observed Confetti expression in the LSK population and T cells (Figure 2A, Figure S2A-E). Unexpectedly, Confetti expression was additionally detected in NK cells, CD41^+^ cells, non-inflammatory monocytes (SSC-A^low^Ly6C^-^Ly6G^-^CD11b^+^), common myeloid progenitors (CMP, Lin^-^ Sca-1^-^cKit^+^CD16/32^-/low^CD34^+^), and granulocyte-monocyte progenitors (GMP, Lin^-^Sca-1^-^ cKit^+^CD16/32^+^CD34^+^) [21] (Figure S2D-G). Considering the relatively shorter lifespan of these cells compared to HSC/MPP, the labeling in these cells should minimally impact precursor calculations, especially with a chase period post-labeling. While the Confetti induction in the relatively long-lived T cells precludes estimation based on mature T cells, we conclude that HSC-Scl-CreER^T^ remains a practical choice for labeling active hematopoietic precursors.

**Figure 2.**
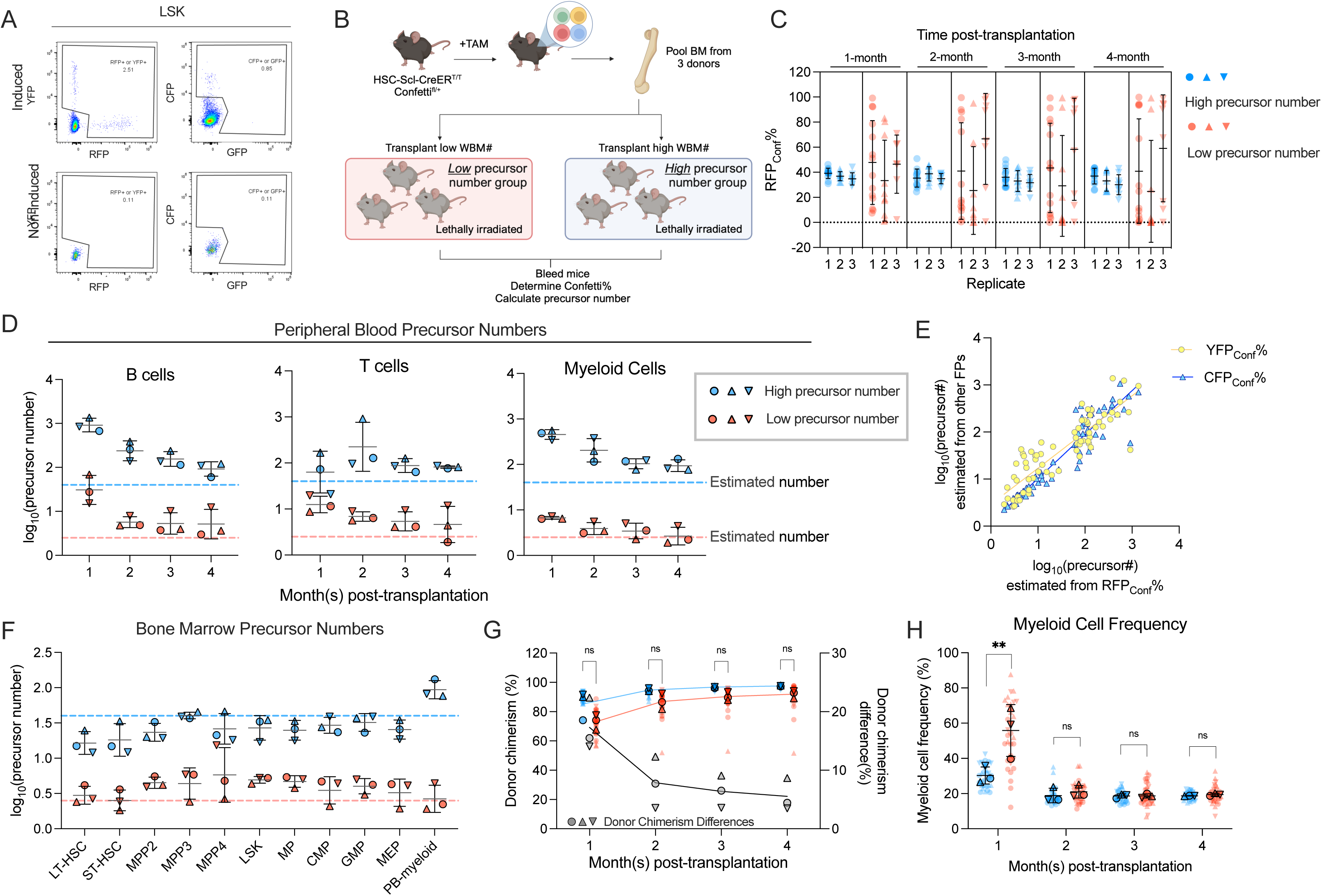
Transplantation of Defined Number of Precursors (A) Representative flow plots showing Confetti induction by HSC-Scl-CreER^T^ in the LSK population. (B) Schematic for transplantation of defined number of precursors. Briefly, 4×10^6^ Confetti-induced WBM for “high precursor number” groups, or 0.25×10^6^ WBM for “low precursor number” groups were transplanted into lethally irradiated recipient mice. Image was created with Biorender. (C) RFP_Conf_% distribution in PB myeloid cells. Each dot represents one animal. Recipient sample size = 7-14 mice per replicate, N = 3 replicates per group. Error bars represent mean ± SD. (D) The precursor numbers of B cells, T cells, and myeloid cells in recipient mice. Dash lines mark the level of historically estimated transplantable clone numbers. Each dot represents a precursor number calculated from multiple mice. Error bars represent mean ± SD. (E) The correlation of precursor numbers calculated from RFP_Conf_%, YFP_Conf_% or CFP_Conf_%. Each dot represents a precursor number calculated from multiple mice. (F) The precursor number of BM MP and HSPC in recipient mice. Dash lines represent historically estimated transplantable clone numbers. Each dot represents a precursor number calculated from multiple mice. Error bars represent mean ± SD. (G) The donor chimerism of recipient mice, and the CD45.2^+^ chimerism differences between high- and low- precursor number groups. Each solid dot represents one animal. Each outlined dot represents the average value of a replicate containing multiple animals. Recipient sample size = 7-14 mice per replicate, N = 3 replicates for each precursor number group. Two-way ANOVA for paired samples was performed. (H) The frequency of myeloid cells in the peripheral blood of recipients. Each dot represents one animal. Recipient sample size = 7-14 mice per replicate, N = 3 replicates for each precursor number group. Two-way ANOVA for paired samples was performed. ns, non-significant, * p < 0.05, ** p < 0.001.

The stability of Confetti labeling after induction is required for binomial distribution. Therefore, the absence of background Cre activity after Confetti induction is critical. We determine the background Cre activity in HSC-Scl-CreER^T^/Confetti animals by two approaches: (1) detecting Confetti expression in non-induced animals; (2) identifying cells co-expressing two Confetti FPs months after tamoxifen induction due to cassette "flipping" from background Cre activity. We observe no Confetti-expressing cells without tamoxifen treatment in mice up to 50 weeks old and minimal co-expression of two Confetti FPs in induced HSC-Scl-CreER^T^/Confetti animals (Figure S3A-B). Conversely, Vav-Cre/Confetti animals constitutively express Cre, resulting in ∼15% cells co-expressing two Confetti colors in peripheral blood (PB) T cells (Figure S3C-D). Both lines of evidence support minimal Cre background activity in HSC-Scl-CreER^T^ and its use as a Cre driver for precursor number calculation.

### The variance of FP% inversely correlates with precursor numbers *in vivo*

To ascertain the inverse correlation between variance of FP% and the number of hematopoietic precursors *in vivo*, we first investigate the variance of FP% in mice hosting various numbers of hematopoietic precursors. We generate these mice by non-competitively transplanting 0.25×10^6^ or 4×10^6^ induced CD45.2^+^ HSC-Scl-CreER^T^/Confetti mouse bone marrow (BM) cells into CD45.1^+^ recipient mice (Figure 2B). Given the constant frequency of precursors in the donor BM (1/10^5^) and the varying doses of donor BM cells, these mice received approximately 2.5 (“low precursor number”) or 40 (“high precursor number”) precursors [16,24–26]. While similar experiments transplanting defined numbers of precursors have been reported, the estimation for precursor numbers lower than 50 has not been explored.

Due to the scarcity of hematopoietic precursors in the BM, the precursor number seeded in recipients followed Poisson distribution instead of being constant, violating the premise (1) of binomial distribution. Nonetheless, our simulations show that the seeding number variation among recipient mice is relatively small, therefore the variance resulting from precursor number differences still dominates and inversely correlates with precursor numbers. A minor problem is that the estimated numbers are not at one-to-one ratio to expected numbers at low precursor number range (<10, see Methods) (Figure S4A).

As cells are not 100% labeled with Confetti, for clarity, we use “FP_PB_%” or “FP_BM_%” to represent the frequency of a given FP in the PB or BM, and use “FP_Conf_%” to represent the frequency of a given FP in the Confetti^+^ population. For total precursor number calculations, we normalize the number calculated from the Confetti-labeled population to Confetti labeling efficiency.

As expected, we observe higher RFP_Conf_% variance across all PB cell types in mice belonging to the “low precursor number” groups (Figure 2C and Figure S4B-C). Since variance of any FP following a binomial distribution inversely correlates with precursor numbers, we also find higher CFP_Conf_% and YFP_Conf_% variance in the “low precursor number” groups (Figure S4D-E). Following the same principle, we observe higher RFP_PB_%, CFP_PB_%, YFP_PB_% and Confetti% variance in “low precursor number” mice (Figure S4F).

Applying equation 2, we find that the precursor numbers of B cells and myeloid cells are noticeably higher in the first two months than at four months post-transplantation, suggesting transient progenitor contributions at early time points (Figure 2D). At four months post-transplantation, the estimated precursor numbers align with the expected values (in myeloid cells, 94 for “high precursor number” groups, three for “low precursor number” groups, Figure 2D). Again, for any FP following binomial distribution, variance of FP% inversely correlates with precursor numbers and thus can be leveraged to calculate precursor numbers (Methods). Since different FPs measure the same population of precursors, we expect the estimations from different FPs to be similar. Indeed, precursor numbers derived from RFP_Conf_% highly correlate with those estimated from YFP_Conf_% or CFP_Conf_% (R^2^ = 0.885 for CFP% in Confetti^+^, R^2^ = 0.769 for YFP% in Confetti^+^), as well as those from RFP_PB_%, YFP_PB_% and CFP_PB_% (Figure 2E, Figure S4G).

Consistent with PB data, higher variance of RFP_Conf_% is also observed in the BM of “low precursor number” mice (Figure S5A). The precursor numbers estimated from various BM HSPC subpopulations align well with each other and are consistent within the same group (Figure 2F). However, in “high precursor number” groups, estimates from BM HSPCs are lower than those derived from PB myeloid cells (27 for BM HSPC, 94 for PB myeloid cells, Figure 2F). This discrepancy may reflect uneven seeding of precursors to the BM throughout the body after transplantation, and the fact that we only sample a part of the BM (femur, tibia, and pelvis)[27]. In summary, we validate the inverse correlation between variance of FP% and precursor numbers *in vivo*. Moreover, the quantification of precursor numbers is feasible even for precursor numbers outside the empirical correlation range.

### Cell frequency measurements fail to reflect the differences in precursor numbers

Given that transplantation studies are performed non-competitively and the recipients are lethally irradiated, we expect to see minimal differences in donor chimerism between two groups, despite drastic differences in donor precursor numbers. Indeed, although we observe substantially lower donor chimerism in mice belonging to “low precursor number” groups during the first two months post-transplantation, the chimerism differences are very small by four months post-transplantation (5.5±2.8%, Figure 2G). The initial differences in donor precursor numbers do not affect PB cell frequencies, except for the first month post-transplantation, when higher PB myeloid frequencies are observed in the “low precursor number” groups (Figure 2H). This suggests myeloid cell production is enhanced when very few precursors are available after irradiation-mediated injury (Figure 2C).

In the BM, nucleated cell counts, donor chimerism and HSPC frequencies are mostly similar regardless of donor precursor numbers, suggesting that even extremely low numbers of hematopoietic precursors can still effectively repopulate a non-competitive environment (Figure S5B-D). Therefore, in cases where hematopoietic precursors expand or decline without competition, stem cell frequencies are less informative to study precursor activity. Measuring precursor numbers is more meaningful as low number of active precursors may be constrained by compensatory proliferation.

### Thousands of hematopoietic precursors contribute to native hematopoiesis

While the empirical formula, which measures 50-2500 precursors, supports quantification of precursors labeled at fetal stages, it may not be applicable to native hematopoiesis, when the total precursor number may exceed 2500. Having validated the feasibility for measuring precursor numbers *in vivo* by Confetti animals, we next sought to investigate the number of active precursors contributing to native hematopoiesis. In HSC-Scl-CreER^T^/Confetti animals, we opt to label a smaller portion of HSPCs with a two-day treatment, despite the potential to label up to 60% HSPCs with Confetti (CFP+YFP+RFP) by 14-day tamoxifen treatment (Figure 3A, Figure S6A). This precautionary measure aims to mitigate potential toxicity arising from prolonged tamoxifen treatment.

**Figure 3.**
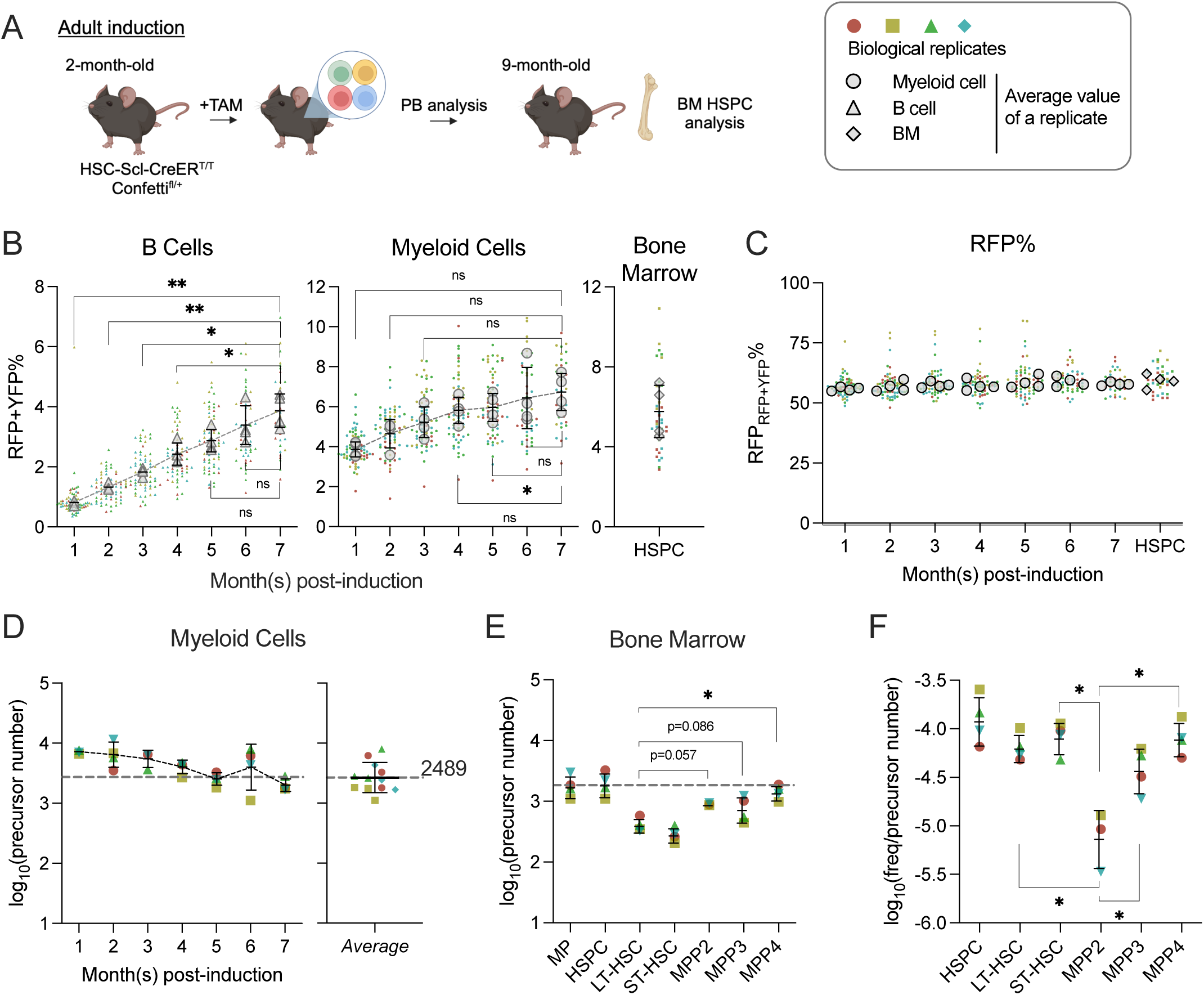
Quantification of active hematopoietic precursor at steady-state (A) Experiment schematic for Confetti induction in adult-induced animals. Image was created with Biorender. (B) Confetti labeling in B cells, myeloid cells and BM HSPCs. Statistics for comparisons between labeling of month seven and other months were shown. One-way ANOVA for paired samples was performed. (C) RFP_Conf_% in PB myeloid cells and BM HSPCs. Statistics for comparisons between labeling of month seven and other months were shown. One-way ANOVA for paired samples was performed. (D) The number of myeloid precursors and the average precursor number from month five to seven. (E) The number of BM precursors. Paired permutation was performed. (F) The frequency-to-clone ratio of bone marrow HSPCs. Paired permutation test was performed. Each solid dot represents one animal. Each outlined dot represents the average value of a replicate containing multiple animals. Error bars represent mean ± SD. PB sample size = 14-22 mice per replicate; BM sample size = 8-10 mice per replicate; N = 4 replicates. ns, non-significant, * p < 0.05, ** p < 0.001.

Unexpectedly, we observed a successive decrease of CFP_Conf_% and CFP_PB_% in some animals, which distort the distribution of CFP_Conf_% (Figure S6B and C). A similar decline is not observed in RFP_Conf_% or YFP_Conf_%, nor in RFP_RFP+YFP_% (calculated by RFP_PB_%/(RFP_PB_%+YFP_PB_%)) (Figure S6D and E). A similar loss of CFP_Conf_% is neither observed in animals with Confetti induction during fetal development, suggesting a potential immune response to CFP in adult-induced animals (Figure S6F). To circumvent this caveat, we decided to calculate precursor numbers solely based on the variance of RFP_RFP+YFP_%. Since only ∼2% of myeloid cells are labeled with CFP, we reason that the decline of CFP in some animals is unlikely to affect the calculation of total precursor numbers based on other FPs, such as RFP_RFP+YFP_%. Moreover, in transplantation studies, precursor numbers calculated with variance of RFP_RFP+YFP_% linearly correlate with those calculated using variance of other FPs (Figure S4G).

To further ensure accuracy, we focus on PB myeloid cells, as (1) Confetti labeling of T cells result from direct induction by HSC-Scl-CreER^T^, but not differentiation from HSPCs (Figure S2D); (2) Confetti labeling (RFP and YFP) of B cells does not plateau at seven months post-induction (Figure 3B); (3) myeloid cells are of shorter lifespan. In myeloid cells, we focus on precursor numbers after four months post-induction, when their labeling reaches a plateau (Figure 3B). The stability of average RFP_RFP+YFP_% suggests an equal contribution of RFP^+^ and YFP^+^ cells to the PB, a prerequisite for estimating precursor numbers from FP% distributions in the progenies (the myeloid cells) (Figure 3C). Like those in PB, the average RFP_RFP+YFP_% in BM HSPC subpopulations is also stable, supporting non-biased differentiation of RFP^+^ and YFP^+^ precursor cells (Figure 3C).

Fitting data to equation 2, we estimate an average of 2667 precursors contributing to native myelopoiesis (Figure 3D, average of numbers at five to seven months post-induction). This number closely aligns with the clone number estimated by transposon-based barcodes statistic in granulocytes (831/30% = 2770) [1].

The average precursor number calculated from PB myeloid cells at the time of BM analysis (1958) matches those calculated from BM myeloid progenitors (MP, Lin^-^Sca-1^-^cKit^+^) and HSPCs (1773 and 1917), but it is five-fold higher than that of LT-HSC (Figure 3E). Although LT-HSCs are thought to sustain steady-state hematopoiesis, recent studies suggest that ST-HSC and MPPs persist long-term as primary contributors to adult hematopoiesis [1,19,21,28]. The observed discrepancy between the number of precursors contributing to LT-HSC and those contributing to MPPs indicates that at least some MPPs are not replaced by progenitors directly differentiated from LT-HSC at seven months post-induction, affirming the persistence of MPPs (Figure 3E). Furthermore, the fact that PB myeloid and BM MP precursor numbers are closer to those of MPPs than LT-HSC confirms active and long-term (at least seven months) contributions from MPPs to steady-state myelopoiesis.

Of note, the quantity of precursor does not necessarily correlate with the frequency of cell type in the BM. For example, the cell frequency of MPP2 (Lineage^-^cKit^+^Sca-1^+^Flk2^-^CD48^+^CD150^+^) in BM is very low, but MPP2 comprises a modest number of precursors, making its frequency-to-precursor-number ratio the lowest among the HSPC subtypes (Figure 3F and S6G). The frequency-to-precursor-number ratio reflects how well precursors of a particular type proliferate to expand their absolute cell count. The low frequency-to-precursor-number ratio of MPP2 suggests that it expands more poorly than other HSPC subtypes.

In summary, we detect thousands of hematopoietic precursors contributing to adult hematopoiesis. At the time of BM analysis, the number of PB myeloid precursors is comparable to those observed in BM MP and HSPCs.

### Precursor numbers determined by FP% variance confirm reduced clonality of progenitors after myeloablation

Myeloablation through 5-fluorouracil (5-FU) treatment depletes most actively cycling cells, forcing quiescent stem cells to proliferate [29]. Previous studies have indicated a significant reduction in the number of clones detected within the BM c-Kit^+^ population (consisting of MP and HSPC) following a single dose of 5-FU treatment [9]. However, questions arise regarding whether this is an artifact stemming from potential under-sampling of the highly expanded c-Kit^+^ population after 5-FU treatment by single-cell sequencing. Given the quantitative estimation of a wide range of precursor numbers through equation 2, we aimed to investigate whether precursor numbers within progenitor populations indeed reduce ten days post-5-FU treatment (Figure 4A).

**Figure 4.**
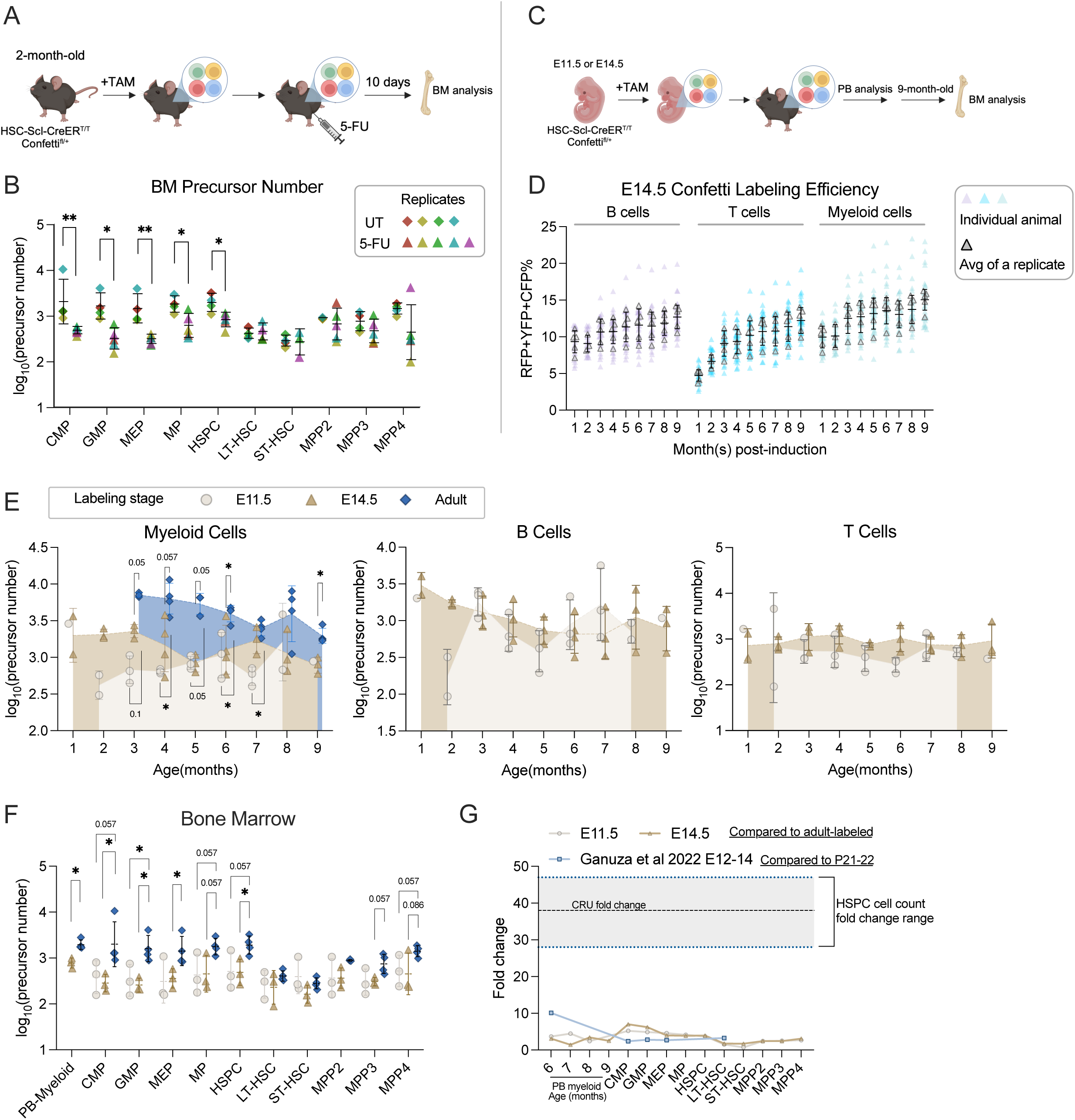
Number of precursor post-5-FU treatment and along developmental ontogeny (A) Experiment schematic for one-dose 5-FU treatment. Image was created with Biorender. (B) The number of BM precursors post-5FU treatment. Permutation test was performed. For UT, sample size = 8-10 mice per replicate; N = 4 replicates; for 5-FU, sample size = 6-8 mice per replicate; N = 5 replicates. (C) Experiment schematic for Confetti induction at fetal stages. Image was created with Biorender. (D) The Confetti labeling of B cells, T cells and myeloid cells. Each dot represents one animal. Each outlined dot represents the average value of a replicate containing multiple animals. Sample size = 4-11 mice per replicate; N = 5 replicates. (E) The number of precursors in PB myeloid cells, B cells and T cells. Permutation test was performed. For E14.5 and adult-induction, N = 4 replicates; for E11.5, n = 3 replicates. (F) The number of BM precursors. Each dots represent one precursor number. Permutation test was performed. For E14.5 and adult-induction, N = 4 replicates; for E11.5, N = 3 replicates. (G) The relative fold change increase of precursor numbers from fetal- to adult-stage, as well as fold change of CRU and cell counts. Error bars represent mean ± SD. ns, non-significant, * p < 0.05, ** p < 0.001.

To determine the precursor changes post-5-FU treatment, we use the animals described in Figure 3A as untreated (UT) benchmark cohort, which were collected at the same age. At ten days post injection, the efficacy of 5-FU treatment is validated by lower PB myeloid cell frequency post-treatment and higher frequencies of BM progenitors compared to UT animals (Figure S7A and B). The high progenitor frequencies in the BM results from the depletion of cycling cells in the BM and the enhanced proliferation of HSPCs following 5-FU treatment.

Compared to UT animals, the precursor numbers of MP (including CMP, GMP, MEP) and HSPC significantly decrease, confirming a reduction in clonality in the c-Kit^+^ population (Figure 4B) [9]. By contrast, the precursor numbers of LT-HSC do not show a decreasing trend. While transplantation studies support unchanged clonality of primitive stem cells after a single dose of 5-FU, a similar investigation in a native environment has not been conducted [30]. Our findings support the notion that native LT-HSC clonality remains unaltered following one-dose 5-FU treatment. Together, myeloablation treatment reinforces how the dynamics of precursor numbers can be tracked through Confetti pattern variations.

### Modest developmental expansion of active lifelong precursors

HSCs are thought to undergo substantial expansion in the fetal liver during fetal development [31–33]. However, a recent study employing the empirical formula challenges this notion by quantifying endogenous lifelong hematopoietic precursors labeled at various developmental stages [34]. It revealed limited expansion of hematopoietic precursors during the fetal liver stage (from E10 to E15,1.8- to 2.7-fold) as well as a gradual and moderate increase from the fetal liver to the post-natal stage (2.4- to 10-fold) [34]. Although intriguing, most post-natal measurements in this study fell outside the empirical linear range, leaving the genuine degree of post-natal precursor expansion uncertain. As our formula quantitatively assesses a wide range of precursor numbers, we set out to directly compare the numbers of precursor labeled at various development stages (E11.5 and E14.5, by one dose of tamoxifen; adult-stage, benchmark shown in in Figure 3A, all analyzed at the same age) (Figure 4C).

For accuracy, we first examine the dynamics of Confetti labeling. In E14.5-induced animals, the Confetti labeling of T cells almost double from one month to four months of age (4.7% at one month, 9.3% at four months), suggesting Confetti^+^-labeled precursors contribute more to post-natal T cells than non-labeled precursors (Figure 4D). Therefore, for T cells in E14.5-induced animals, we focus on precursor numbers generated after three months of age. The Confetti labeling for E11.5-induced animals remain stable for all timepoints examined (Figure S7C).

Similar to adult-induction, the distribution of average FP% is stable for fetal-induction, supporting equal proliferation of RFP^+^ and YFP^+^ cells (Figure S7D). Unlike adult-induction, fetal-labeled precursor numbers can be calculated with various combination of FP%, since the expression of CFP introduced at the fetal stages is stable (Figure S6F). Nonetheless, for consistency, all precursor numbers are calculated with variance of RFP_RFP+YFP_%, the one employed in adult-induction animals.

The resulting precursor number estimates for E11.5 and E14.5-labeled cohorts are similar in all PB cell types except at two and three months of age, echoing the previous study reporting limited lifelong precursor expansion in the fetal liver (Figure 4E) [34]. Precursor numbers calculated from BM subpopulations are also comparable between the two timepoints (Figure 4F). Although E9.5-labeled clones are reported to seed non-uniformly across bones, the precursor numbers calculated from PB myeloid cells do not significantly differ from those calculated from BM MP and HSPC in E14.5-labeled animals (Figure 4F) [9].

For comparison between fetal- and adult-induced animals, we focus on PB myeloid cells, as adult-induced animals have non-saturated labeling in B cells and non-HSPC-rooted Confetti labeling in T cells (Figure 3B). Here, we observe a relatively small increase of precursor numbers in adult-induced animals in PB and BM compared to those labeled at the fetal stage (Figure 4E and F). The increase between E11.5/E14.5- and adult-labeled precursors is less than ten-fold, similar to a previous report (Figure 4G) [34].

Together, we confirm minimal to no expansion of lifelong precursors in the fetal liver stage and a minor expansion of expansion of lifelong precursors from fetal liver to adult stage.

### *Fancc^-/-^* mice have normal numbers of hematopoietic precursors at steady-state

Our approach provides an opportunity to investigate the number of active HSPC precursors in the native environment. To showcase precursor quantification in genetic mouse models, we focus on Fanconi Anemia (FA), the most common inherited bone marrow failure syndrome [35]. Current FA mouse models exhibit mostly normal adult hematopoiesis at steady state but demonstrate reduced repopulation ability upon BM transplantation [36–41]. It remains unclear if their steady-state blood production is sustained by a reduced number of precursors, predisposing them to repopulation defects after transplantation.

To quantify precursor number in a mouse model of FA, we generate *Fancc*^+/+^, *Fancc*^+/-^ and *Fancc*^-/-^ mice with a single Confetti allele and homozygous HSC-Scl-CreER^T^ alleles (Fancc^+/+^Confetti^fl/+^HSC-Scl-CreER^T/T^, hereafter *Fancc*^+/+^; Fancc^+/-^Confetti^fl/+^HSC-Scl-CreER^T/T^, hereafter *Fancc*^+/-^; Fancc^-/-^ Confetti^fl/+^HSC-Scl-CreER^T/T^, hereafter *Fancc*^-/-^) [42]. Consistent with previous literature, we observe similar PB cell frequencies and blood counts between *Fancc*^+/+^ and *Fancc*^-/-^ mice (Figure S8A-C) [41]. BM nucleated cell counts, as well as HSPC and MP frequencies, are mostly identical, except for ST-HSC frequencies, which are significantly reduced in *Fancc*^-/-^ mice compared to *Fancc*^+/-^ mice (Figure S8D-E).

To label hematopoietic precursors in FA animals, Confetti expression was induced at two months of age, and FP% was monitored over seven months (Figure 5A). The absence of *Fancc* did not affect Confetti labeling efficiency, Confetti labeling dynamics or average FP% (RFP_RFP+YFP_%), suggesting precursor numbers in *Fancc^-/-^* can be similarly calculated (Figure 5B and Figure S8F). As in previous adult-induction animals, RFP_RFP+YFP_% was used to estimate precursor numbers. The resulting myeloid and BM precursor numbers were comparable regardless of *Fancc* genotype, albeit *Fancc*^-/-^ mice had a slight reduction in HSPC precursor numbers (Figure 5C and D). These observations collectively suggest that a normal number of precursors sustains blood production in *Fancc^-/-^* mice.

**Figure 5.**
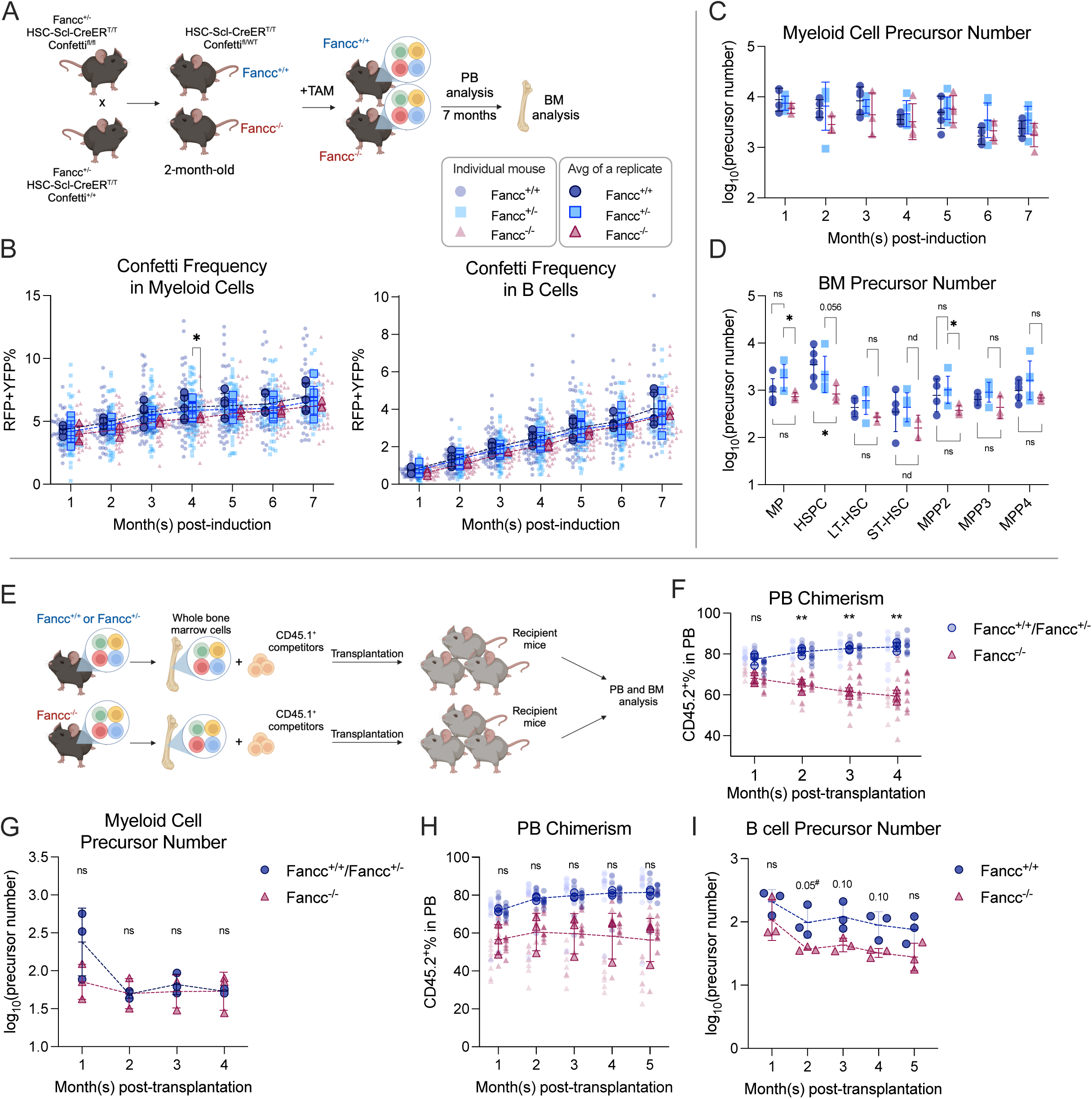
Precursor numbers in a mouse model of FA (A) Experimental workflow to generate *Fancc*^+/+^ and *Fancc*^-/-^ mice, induce Confetti expression and track precursor number dynamics. Image was created with Biorender. (B) Confetti labeling in PB B cells and myeloid cells. Each dot represents one animal. Each outlined dot represents the average value of a replicate containing multiple animals. Dashed lines connect the values for the same replicate. (C) The number of PB myeloid precursor. (D) The number of BM precursor. Permutation test was performed. (E) Experiment schematic for competitive transplantation. Image was created with Biorender. (F) PB donor chimerism in recipient mice of young FA donors. Each dot represents one animal. Dashed lines connect the average donor chimerism of three replicates. For Fancc^+/+^ or Fancc^+/-^, n = 10 per replicate, N = 3 replicates; for Fancc^-/-^, N = 7-9 per replicate, N = 3 replicates. Two-way ANOVA was performed. (G) The number of PB myeloid precursor post-transplantation of young FA donors. Dashed lines connect the average precursor numbers. Permutation test was performed. N = 3 for each group. (H) PB donor chimerism recipient mice of aging FA donor cells. Each dot represents one animal. Dashed lines connect the average donor chimerism of three replicates. For both genotype, N = 10 per replicate, N = 3 replicates. Two-way ANOVA was performed. (I) The number of PB B precursor post-transplantation of aging FA donors. Dashed lines connect the average precursor numbers. Permutation test was performed. N = 3 for each genotype. For (A-C), Fancc^+/+^, sample size = 11-17 mice per replicate, N =5 replicates; Fancc^+/-^, sample size = 10-22 mice per replicate, N =7 replicates; Fancc^-/-^, sample size = 9-17 mice per replicate, N =4 replicates. For (D), Fancc^+/+^, sample size = 5-9 mice per replicate, N =5 replicates; Fancc^+/-^, sample size = 6-12 mice per replicate, N =5 replicates; Fancc^-/-^, sample size = 6-11 mice per replicate, N =4 replicates. Error bars represent mean ± SD. nd, not determined; ns, non-significant; * p < 0.05, ** p < 0.001. # represents lowest p value possible for permutation test.

### The number of *Fancc^-/-^*precursors remains unchanged in mice transplanted with young donor cells

While *Fancc*^-/-^ mice have a similar number of precursors as their wildtype counterparts at homeostasis, it is unknown whether the reduced repopulation ability post-transplantation stems from diminished precursor numbers, reduced cell expansion, or a combination of both [37]. To understand the underlying mechanism, we performed competitive transplantation using BM cells from three-month-old Confetti-induced *Fancc*^+/+^, *Fancc*^+/-^ or *Fancc*^-/-^ mice along with CD45.1^+^ competitor cells (Figure 5E, Figure S9A). Consistent with previous studies, recipient mice of *Fancc*^-/-^ cells show significantly lower donor chimerism in the PB and BM compared to those of *Fancc*^+/+^ or *Fancc*^+/-^ cells (Figure 5F, Figure S9B). Despite lower donor chimerism, PB and BM precursor numbers were unaffected in *Fancc*^-/-^ recipient mice (Figure 5G, Figure S9C-E). This suggests that reduced *Fancc*^-/-^ cell proliferation instead of fewer active precursors is likely the cause for the reduced repopulation capacity post-transplantation.

### Ageing *Fancc*^-/-^ mice have reduced lymphoid hematopoietic precursors upon transplantation

Ageing *Fancc*^-/-^ mice have been reported to develop hematologic neoplasms, resulting in decreased survival [43]. To determine if aging *Fancc^-/-^* cells also maintain similar precursor numbers post-transplantation, we competitively transplanted BM cells from nine-month-old Confetti-induced *Fancc^+/+^* or *Fancc^-/-^* mice with CD45.1^+^ competitor cells, a stage when aging *Fancc^-/-^*mice start to show decreased survival [43] (Figure S9F).

The diminished repopulation ability of *Fancc^-/-^* cells is reaffirmed by lower PB and BM chimerism (Figure 5H, Figure S9I). While no differences in precursor numbers are noted during the initial post-transplantation period, a slight yet consistent reduction in lymphoid precursors is observed in *Fancc^-/-^*recipient mice at three to five months post-transplantation (Figure 5I, Figure S9G and H). Changes in precursor numbers in the BM of KO recipients are less clear, as we observed high variance in several HSPC subtypes (Figure S9J). Nonetheless, precursor numbers of MEP, LSK and ST-HSC showed a consistent reduction. In conclusion, aging *Fancc^-/-^* mice showed a modest but consistent loss of active PB lymphoid precursors post-transplantation, implying decreased lymphoid precursors additionally compromise repopulation capacity as *Fancc^-/-^* mice age.

## Discussion

The polyclonal nature of endogenous hematopoiesis imposes methodological problems on a robust dynamic measurement. Inspired by the XCI studies, we successfully employed the correlation formula informed by binominal distribution to quantify precursor number in native hematopoiesis. Using HSC-Scl-CreER^T^/Confetti animals, we estimate thousands of precursors contributing to adult native hematopoiesis, a number comparable to the previous report [1]. This number results from a moderate increase during fetal-to-adult transition and respond dynamically to myeloablation by 5-FU. Applied to a mouse model of inherited bone marrow failure (Fancc^-/-^ mice), we detected normal precursor numbers at steady state and decreased lymphoid precursors upon transplantation of aging donors.

Although the linear relationship between variance of FP% and precursor numbers has been described, its measurable range has been limited [11]. Estimates outside this linear range may erroneously fall within it, making it challenging to distinguish accurate measurements within the range from inaccurate ones beyond it [11]. Based on binomial distribution, we expanded the measurable range to encompass the full spectrum of hematopoietic precursors in mice, minimizing the likelihood of inaccuracies. Furthermore, the linear correlation based on binomial distribution can accommodate any labeling system that follows the underlying premises, without the need for the multispectral Confetti cassette.

Having established a quantitative measurement, we were initially surprised by the moderate differences between fetal and adult-precursors [34]. The maximum increase observed (seven-fold observed in CMP) was substantially lower than the increase of competitive repopulation units (CRU) from E12 to E16 determined by limited dilution assays (∼38-fold) (Figure 4G), as well as the escalation from immunophenotype-defined HSPC counts (28- to 47-fold, from 3-5×10^3^ at E14.5 to 1.4×10^5^ at two months old) (Figure 4G) [18,32,44,45]. It is possible that we overestimated E11.5 precursors, as the numbers of lifelong E11.5 precursors were substantially higher (on average 753 PB myeloid precursor, 512 HSPC precursors) than the one to two repopulation units estimated with transplantation [46]. Nonetheless, our results were comparable to the ∼870 hematopoietic cluster cells observed at E11.5 [47]. We may have underestimated precursor numbers in adult-labeled animals, as most adult HSCs are quiescent [29]. However, HSC precursor numbers did not increase after induced proliferation by one dose 5-FU treatment (Figure 4B). Therefore, we confirm the moderate precursor numbers difference between fetal- and adult-stage, emphasizing the analysis of hematopoiesis in a native environment. Future studies using different methods to investigate precursor activity locally should validate this result.

Novel methods tracing clones *in situ* offer new opportunities to study native hematopoiesis, yet most are challenging to apply in genetic models [1,9,10,27,48]. Since only two mouse lines are required (Confetti and Cre, sometimes Cre is already included for conditional deletion of alleles), our approach is particularly convenient to study hematopoiesis in mouse models of genetic disorders. Applying precursor measurements to *Fancc^-/-^* mice, we observe a minor decrease in precursor numbers after transplantation of aging donors. Although donor chimerism differences had been linked to differences in precursor numbers, reduced proliferation capacity may also contribute to reduced competitive repopulation capacity [26]. In this case, definitive precursor number analysis is necessary to differentiate these possibilities.

Currently, concerns regarding clonal restriction in the context of FA gene therapy arises, as the engraftment of gene-corrected stem cells has yielded marginal clinical benefits [49]. For the first time, we demonstrate that FA cells maintain a normal precursor quantity post-transplantation, thereby disproving clonal attrition, including putative homing deficits post-transplantation as a cause in this model [50]. However, it is imperative to acknowledge that the majority of murine models of FA, including the *Fancc^-/-^*model utilized herein, do not recapitulate the pathophysiology observed in FA patients. Future investigations in other FA mice and studies leveraging FA patient-derived materials will be pivotal in corroborating and validating the findings presented here.

While we implemented a careful data processing procedure, one pitfall of our analyses was that we inferred precursor numbers from their progenies, assuming uniform and linear expansion from precursor to progenies. A recent study showed non-uniform precursor clone sizes, although the level of non-uniformity is low [27]. In cases where the non-uniformity is high, according to mathematical modeling, we measured the major contributors to hematopoiesis[51]. Another potential caveat is that the relative contribution of Confetti-labeled precursors to blood production compared to non-labeled precursors remains unknown. Future studies using different Cre drivers should validate the precursor numbers for steady-state hematopoiesis.

In summary, we substantially broadened the applicable range of the correlation between variance of FP% and the precursor based on binomial distribution. We discovered thousands of precursors contributing to steady-state adult murine hematopoiesis and validated that fetal-to-adult precursor expansion is indeed limited. This analysis highlights active precursor numbers as an important metric in both normal and genetic mouse models.

## Methods

### Variance modeling with two-color HL-60

HL-60 cells were cultured with IMDM containing 20% FBS (Gemini) and 5% Penicillin-Streptomycin (Gibco) and maintained at 1 x 10^5^ and 1 x 10^6^ cells/ml. Mycoplasma tests (Lonza) were performed routinely to rule out mycoplasma contamination. BFP-HL-60 and GFP-HL-60 were generated with lentivirus transduction of pGK-BFP (Genscript) and LeGO-V2 (a gift from Dr. Stefano Rivella lab). For seeding of one to 10,000 cells, sorting of HL-60 mixtures was performed using a BD FACS Aria III. For seeding of 100,000 cells, cell counts and dilution was used. After expansion, HL-60 cells were fixed with BD fixation buffer before Aurora (Cytek) or Cytoflex (Beckman Coulter) analysis.

### Mice

HSC-Scl-CreER^T^ mice [23] were crossed with R26R-Confetti mice (B6.129P2-*Gt(ROSA)26Sor^tm1(CAG-Brainbow2.1)Cle^*/J) to generate HSC-Scl-CreER^T/T^Confetti^fl/+^ (HSC-Scl-CreER^T^/Confetti) animals. HSC-Scl-CreER^T^/Confetti animals were crossed with *Fancc*^+/-^ mice [42] to generate HSC-Scl-CreER^T/T^Confetti^fl/+^*Fancc*^+/+^ (*Fancc^+/+^*), HSC-Scl-CreER^T/T^Confetti^fl/+^*Fancc*^+/-^ (*Fancc^+/-^*), and HSC-Scl-CreER^T/T^Confetti^fl/+^Fancc^-/-^ (*Fancc^-/-^*) mice. Vav-Cre was generously offered by Dr. Wei Tong (Children’s Hospital of Philadelphia). Vav-Cre/Confetti animals were generated by crossing Vav-Cre with R26R-Confetti mice. All animals are in the B6 (CD45.2) strain background, unless otherwise stated. Six- to twelve-week-old females were used for timed pregnancies. Eight-week-old mice were used for adult Confetti induction. Both female and male mice were used [11]. Six- to eight-week-old old female B6 CD45.1 mice (B6.SJL-*Ptprc^a^ Pepc^b^*/BoyJ, Jackson laboratories) were used as competitors and recipients for transplantation studies. All mice were maintained in the conventional small animal facility at the Children’s Hospital of Philadelphia (CHOP). All procedures involving animals were approved by the Institutional Animal Care and Use Committee at the Children’s Hospital of Philadelphia.

### Animal identification

Tail snip DNA was extracted using KAPA Express Extract Kit (Roche). Genotyping PCR was performed with HotStarTaq Master Mix (Qiagen) according to manufacturer’s instruction. Genotyping primers used are summarized in Table S3. To determine the zygosity of HSC-Scl-CreER^T^, qPCR was additionally performed with purified tail snip DNA using SYBR™ Green Universal Master Mix (Applied Biosystems).

### Animal procedures

For fetal induction, timed matings of HSC-Scl-CreER^T/T^Confetti^fl/fl^ mice and HSC-Scl-CreER^T/T^ mice were set up. The mice were separated the next morning and noon of the day of separation was considered E0.5. Tamoxifen was delivered at 100mg/kg to the dam orally at E11.5 or E14.5. Pups were C-sectioned and cross-fostered at E18.5 due to reported delivery difficulties caused by tamoxifen treatment [52]. For mice used for defined (“low” versus “high”) number of transplantation (Figure 2), tamoxifen was delivered at 70mg/kg orally once per day for 14 days at eight weeks old. For adult induction (Figure 3), tamoxifen was delivered at 70mg/kg orally once per day for two days at eight-week-old. For one dose 5-fluorouracil (5-FU, Sigma) treatment, 5-FU was intraperitoneally injected once ten days before BM harvest (37-week-old) at 150 mg/kg. To obtain peripheral blood for Confetti analysis, mice were anesthetized using isoflurane and retro-orbitally bled or submandibularly bled for *Fancc^-/-^* mice with occasional congenital eye defects[53] for 1 capillary of blood (50ul). For peripheral blood counts, blood was collected in EDTA tubes using similar bleeding methods and was analyzed by the Translational Core Lab (Children’s Hospital of Philadelphia).

### Bone marrow sample processing

To collect BM cells, mice were euthanized by CO_2_ inhalation. Tibia, femur and pelvis were dissected, and the bone marrow cells were flushed with 26-gauge needles. The single-cell suspension generated with 18-gauge needles was then filtered through 70µm strainers. Erythrocytes in the bone marrow cells were hemolyzed by RBC lysis buffer before antibody staining or transplantation. For stem cell enrichment, bone marrow cells were further lineage depleted using EasySep™ Mouse Hematopoietic Progenitor Cell Isolation Kit (STEMCELL Technologies).

### Flow cytometry analysis

Peripheral blood and bone marrow cells were analyzed using Aurora (Cytek) or BD FACS Aria III and the flow cytometry data were analyzed using FlowJo (Tree Star). The combinations of the following cell surface markers were used to define the peripheral blood populations: myeloid cells: CD11b^+^ or Gr-1+; T-cell: CD3ε^+^; B-cell: B220^+^. The following combinations of cell surface markers were used to define the bone marrow stem and progenitor cells (Lineage/Lin: CD11b, Gr-1, B220, CD3ε, Ter119): LTHSC: Lin^-^c-Kit^+^Sca1^+^Flk2^-^CD150^+^CD48^-^; MPP2: Lin^-^c-Kit^+^Sca1^+^Flk2^-^ CD150^+^CD48^+;^ MPP3: Lin^-^c-Kit^+^Sca1^+^Flk2^-^CD150^-^CD48^+^; MPP4: Lin^-^c-Kit^+^Sca1^+^Flk2^+^CD150^-^CD48^+^; STHSC: Lin^-^c-Kit^+^Sca1^+^Flk2^-^CD150^-^CD48^-^; MEP: Lin^-^c-Kit^+^Sca1^-^CD34^-^CD16/32^-^; CMP: Lin^-^c-Kit^+^Sca1^-^CD34^mid^CD16/32^mid^; GMP: Lin^-^c-Kit^+^Sca1^-^CD34^+^CD16/32^+^. For bone marrow stem and progenitor cell analysis, DAPI (Biolegend) was used to distinguish dead cells. Representative examples of flow cytometry gating can be found in Figure S2C. The antibodies were used at optimized dilutions listed in Table S1.

### Bone marrow transplantation

The day before transplantation, female CD45.1 recipient mice were lethally irradiated (5.2Gy x 2, 3h apart) using an X-ray irradiator (Precision). On the day of transplantation, bone marrow cells from donor mice (CD45.2^+^) were collected under sterile conditions as described, RBC-lysed and counted for cell number. For Figure 2 (transplantation of defined precursor numbers), donor mice were induced to express Confetti as described; donor bone marrow cells were (non-competitively) injected into irradiated recipient mice; each matched high- and low- precursor number group received donor bone marrow cells pooled from three to five mice. For transplantation of young *Fancc* mice, donor mice were induced to express Confetti at E14.5; 1.5 x 10^6^ donor bone marrow cells were mixed with 2.5 x 10^5^ CD45.1 supporting bone marrow cells and injected into irradiated recipient mice via tail vein. For transplantation of aging Fancc mice, donor mice were induced to express Confetti at two-months of age; 2 x 10^6^ donor bone marrow cells were mixed with 5 x 10^5^ CD45.1 supporting bone marrow cells.

### Derivation of the correlation between variance of confetti FP% and precursor numbers based on binomial distribution

When a random variable adheres to binomial distribution, studies have established that the following equation holds true:

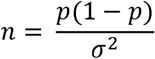

where *n* signifies the number of precursors, *p* represents the probability of an individual being one of the FPs (e.g., RFP), and σ^2^ denotes the variance of specific FP% (e.g. σ_RFP%_^2^)[7]. In experiments, *p* will be estimated with the average FP% in the Confetti^+^ cells (e.g. RFP_mean_%), and σ^2^ will be estimated with the variance of FP% among a group of individual or mice (e.g. 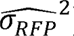). Consequently, the estimation of precursors number n is calculated using the following equation:

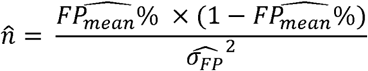

A logarithmic transformation can then be performed, allowing us to establish a linear relationship between the variance of FP% and the number of precursors:

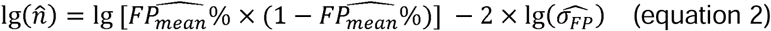

In a previous study[11],

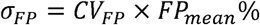

Therefore,

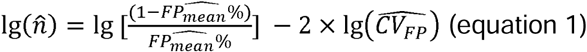

### Resampling to determine sample size per replicate

FP% data generated from HL-60 was used for re-sampling. For each seeded number, FP% was resampled for different sample sizes from all the FP% with replacement. Variance of estimated n was then calculated by the standard deviation of the estimated precursor numbers generated from the same sample size. Relative error was determined by dividing variance of estimated n with the average of the estimated n. Refer to “varying well numbers.rmd” for detailed R code.

### Simulation to determine the effect of varying FP_mean_% on correlation between variance of FP% and precursor numbers

Simulation of binomial distribution of varying FP_mean_% (probability of being a FP) was performed in R, generating corresponding FP% values used for variance calculation. The correlation between variance and precursor number n was then determined by linear correlation, and the slopes and intercepts were compared to each other. Refer to “Simulation of varying FPmean_percent.rmd” for detailed R code.

### Simulation to determine the effect of precursor numbers following Poisson post-transplantation

Precursor numbers in individual samples were simulated to follow a Poisson distribution, where the mean of precursor numbers is the expected precursor numbers (expected n). The induction of a FP in each precursor was then simulated by random assignment, where probability of being a FP was set to be FP_mean_%. For each expected precursor numbers, variance of FP% among samples was then calculated, and equation 2 was used to estimate precursor numbers (estimated n). The correlation between expected n and estimated n was then plotted. Refer to “Double layer binomial simulation.rmd” for detailed R code.

### Data processing and normalization

Peripheral blood cell subset frequencies were normalized to the total % of myeloid cells, T-cell and B-cell, to avoid an underestimate due to incomplete RBC lysis. For transplant animals, peripheral blood CD45.1^+^% and CD45.2^+^% are normalized to total CD45.1^+^% and CD45.2^+^% to avoid an underestimate due to incomplete RBC lysis. After normalization of cell frequencies, the sum of Confetti% and FP_Conf_% (diving FP_PB_% or FP_BM_% by sum of Confetti%) were then calculated for each cohort. At each step, potential outliers were removed based on Tukey method. The variance of FP_Conf_% and average FP_Conf_% are then fitted into equation 2 to calculate precursor numbers if sample size is at least five. The precursor number is then used to compared with the minimum flow cytometry recorded event of samples. If the estimated number is higher than the minimum flow count of samples, the sample with the minimum flow count will be excluded, and the calculation is performed again on the rest of samples. After exclusion, the resulting precursor number will be compared again with minimum flow count of samples, until it is lower than the minimum flow count or sample size is smaller than five.

### Statistical analysis

#### Statistical significance for precursor number estimates

As the distribution of precursor numbers is not predetermined, to compare mean precursor number differences between two conditions from a limited number of biological replicates, we employed permutation test. For unpaired permutation test, if there are three to five biological replicates per condition, the smallest p values possible are 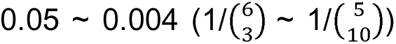. For paired permutation test, if there are three to five biological replicates per condition, the smallest p value possible are 0.125 ∼ 0.03125 (1/2^3^ ∼ 1/2^5^). Thus, for three to five biological replicates per condition, even though some of the p values may not reach the commonly used alpha level (0.05), it still represents substantial number differences. For those p values that were lowest possible but did not reach the alpha level, we specifically labeled with a “#” in the figures and legends.

### Statistical analyses of other data

All other two-sample statistical analyses were performed using Student’s t test, if the sample was normally distributed, or Welch’s t test, when the sample was not normally distributed (F test). For multiple comparison, one-way ANOVA or two-way ANOVA was used.

## Data and code availability

The original Confetti% measured in the peripheral blood and the bone marrow and R code to analyze and reproduce all the results, numeric and figures can be found at https://doi.org/10.5281/zenodo.8222789. The re-analyzed data from a previous study can be found in the online version of papers[11].

## Acknowledgments

Work was supported by R01-HL150882. We thank Florin Tuluc (CHOP Flow Cytometry Core) and Jessica Gucwa (Cytek bioscience) for assistance in flow cytometry; Kaosheng Lv (CHOP) for preparation of reagents; Zilu Zhou (Penn) for help in statistical analysis; Nancy Speck (Penn), Wei Tong (Penn), Julia Warren (Penn), Ding-wen (Roger) Chen (CHOP), Stephanie N Hurwitz (Penn), Hua Qing (Genentech), Hui Chen (Penn) for in-depth discussion. Some figure panels were generated using Biorender.

## Contributions

S.L designed, performed experiments, analyzed data and wrote the manuscript. S.A, J.E, S.J, G.J and T.M performed experiments. H.Z and H.L designed statistical analysis. L.P contributed reagents, guidance and edited the manuscript. P.K conceived and supervised the study and edited the manuscript.

## Supplementary Figure Legend

Figure S1: Establishing the correlation between variance of FP and precursor numbers.

(A) The correlation between precursor numbers and CVs of individual FP or all CVs (original correlation formula), replotted from Ganuza et al., 2017.
(B) FP_mean_% in a previously published dataset, replotted from Ganuza et al., 2017.
(C) The correlation between precursor numbers and standard deviation (σ) of individual FP, replotted from Ganuza et al., 2017.
(D) The BFP% in HL-60 mix during two weeks of growth.
(E) The effect of outliers on precursor number estimation. Left, BFP% in samples with/without outliers. Quartiles by Tukey method was shown. The outlier is highlighted by circle. Middle, standard deviation of BFP% with/without outliers. Right, estimated precursor numbers with/without outliers.
(F) The changes of standard deviation when recorded events varied. “&” denotes the potential overestimated standard deviation due to low flow recorded events.
(G) The variance of estimated precursor numbers when varying sample sizes were used.
(H) The relative error of estimation as sample sizes increased. Each dot represents data from one seeded number. Error bars represent mean ± SD.

Figure S2: The induction specificity of HSC-Scl-CreER^T^

(A) Experiment schematic to examine the induction specificity of HSC-Scl-CreER^T^. N =10 for induced animals, N =3 for non-induced animals. Image was created with Biorender.
(B) Flow gating strategy for BM HSPC and MPs.
(C) Confetti labeling in BM HSPCs. Error bars represent mean ± SD.
(D) Flow gating strategy for BM lineage cells.
(E) Confetti labeling in BM lineage cells. Error bars represent mean ± SD.
(F) Flow gating strategy to examine spleen cells.
(G) Confetti labeling in spleen cells. Error bars represent mean ± SD.

See Methods and Table S1 for the detailed antibody, fluorophore and marker combination used for each cell type.

Figure S3: Analysis of HSC-Scl-CreER^T^ background Cre activity

(A) Representative flow plot showing Confetti expression in PB T cells in 48-50-week-old HSC-Scl-CreER^T^/Confetti animals compared to an animal induced with one-day tamoxifen treatment.
(B) Quantification of RFP and YFP% in PB T cells. N =8 for non-induced 48-50-week-old animals.
(C) Representative flow plot for Confetti labeling in PB T cells post HSC-Scl-CreER^T^-mediated induction.
(D) The level of CFP^+^ and RFP^+^ double positive cells in PB T cells of animals induced by HSC-Scl-CreER^T^ compared to Vav-Cre/Confetti animals analyzed at the same age. N =3 for Vav-Cre animals, N= 8 for HSC-Scl-CreER^T^ animals.

Figure S4: Variance of FP% inversely correlates with precursor numbers *in vivo*

(A) The correlation between expected numbers and the estimated numbers calculated from FP% of precursors following Poisson distribution. See Methods for simulation details.
(B) Flow gating strategy for PB B cell, T cell and myeloid cell. See Methods and Table S1 for the detailed antibody, fluorophore and marker combination used for each cell type.
(C) RFP_Conf_% distribution in B cell and T cell.
(D) YFP_Conf_% distribution in myeloid cell.
(E) CFP_Conf_% distribution in myeloid cell.
(F) RFP_PB_%, CFP_PB_%, YFP_PB_% and Confetti% distribution in myeloid cells.
(G) Linear correlation of PB precursor numbers calculated from RFP_Conf_%, RFP_PB_%, CFP_PB_%, YFP_PB_%, Confetti% or RFP_RFP+YFP_%, Each dot represents a precursor number calculated from multiple mice.

For (C-F), each dot represents one animal. Recipient sample size = 7-14 mice per replicate, N =3 replicates per group. Error bars represent mean ± SD.

Figure S5 BM analysis for high- and low-precursor number groups

(A) RFP_Conf_% distribution in BM HSPCs. Each dot represents one animal
(B) BM donor chimerism.
(C) BM nucleated cell numbers. N =15 mice for high precursor number group, n= 14 for low precursor number group. Unpaired t-test was performed.
(D) BM HSPC and MP frequency in donor CD45.2^+^ population. Two-way ANOVA for paired samples was performed.

For (B) and (D), each solid dot represents one animal; each outlined dot represents the average value of a replicate containing multiple animals. Except for (C), recipient sample size = 7-14 mice per replicate, N =3 replicates per group. For all graphs, Error bars represent mean ± SD. ns, non-significant, * p < 0.05.

Figure S6 Analysis of FP induction in adult-induced animals

(A) Confetti labeling in LSK after 14days of tamoxifen treatment. N =5 animals. Error bars represent mean ± SD.
(B) CFP_Conf_% in PB myeloid cells.
(C) CFP_PB_% in PB myeloid cells.
(D) RFP_RFP+YFP_% in PB myeloid cells.
(E) RFP_PB_% and YFP_PB_% in PB myeloid cells.
(F) CFP_Conf_% in PB myeloid cells of fetal-induced animals
(G) BM HSPC frequencies. Error bars represent mean ± SD.

For (B-E) sample size = 14-22 mice per replicate; for (F), sample size = 4-11 mice per replicate; for (G) sample size = 8-10 mice per replicate. For (B-E) and (G), N =4 replicates. For (F), N =5 replicates.

Figure S7: Analysis related to 5-FU treatment and Confetti fetal induction

(A) PB myeloid cells frequencies before 5-FU treatment and ten days post-5-FU treatment. Dashed lines connect the values for the same replicate. Sample size = 6-8 mice per replicate, N =5 replicates for 5-FU. Paired t test was performed.
(B) The BM HSPC frequencies of UT and 5-FU treated groups ten days post-5-FU treatment. For UT, sample size = 8-10 mice per replicate, N =4 replicates; for 5-FU, sample size = 6-8 mice per replicate, N =5 replicates.
(C) Confetti% in PB in E11.5-labeled animals. Dashed lines connect values for the same replicate.
(D) The distribution of FPs in Confetti^+^ population of B cells. Data showing average from all mice. N =24-34 for E14.5, N =7-24 for E11.5. Error bars represent mean ± SD.

For (A-C), each solid dot represents one animal; each outlined dot represents the average value of a replicate containing multiple animals. ns, non-significant; * p < 0.05, ** p < 0.001.

Figure S8 Analysis of FA mice at steady-state

(A) PB myeloid cell frequencies. Each dot represents one animal. Each outlined dot represents the average value of a replicate containing multiple animals.
(B) PB B cell frequencies. Each dot represents one animal. Each outlined dot represents the average value of a replicate containing multiple animals.
(C) Complete blood count analysis. Each dot represents one animal. N =13 for *Fancc*^+/+^, N =16 for *Fancc*^-/-^ mice. Unpaired t test was performed.
(D) BM HSPC frequencies. Each dot represents one animal. Each outlined dot represents the average value of a replicate containing multiple animals. One-way ANOVA was performed.
(E) BM nucleated cells counts. Each dot represents one animal. Unpaired t test was performed. N =12 for *Fancc*^+/+^and *Fancc*^-/-^ mice.
(F) RFP_RFP+YFP_% in PB myeloid cells and B cells. Each dot represents one animal. Each outlined dot represents the average value of a replicate containing multiple animals.

For (A-B) and (F), Fancc^+/+^, sample size = 11-17 mice per replicate, N =5 replicates; Fancc^+/-^, sample size = 10-22 mice per replicate, N =7 replicates; Fancc^-/-^, sample size = 9-17 mice per replicate, N =4 replicates. For (D), Fancc^+/+^, sample size = 5-9 mice per replicate, N =5 replicates; Fancc^+/-^, sample size = 6-12 mice per replicate, N =5 replicates; Fancc^-/-^, sample size = 6-11 mice per replicate, N =4 replicates. Error bars represent mean ± SD. ns, non-significant; * p < 0.05.

Figure S9 Analysis of FA recipient mice

(A) Young *Fancc* donor BM HSPC frequency analysis. N = 5 for Fancc^+/+^ or Fancc^+/-^, N = 2 for Fancc^-/-^.
(B) Donor chimerism in recipient BM of young *Fancc* donor. Each dot represents one animal. Each outlined dot represents the average value of a replicate containing multiple animals. Two-way ANOVA was performed. Recipient sample size = 7-10 mice per replicate, N =3 replicates per group.
(C) Number of PB B cell precursor in recipients of young FA donor.
(D) Number of PB T cell precursor in recipients of young FA donor.
(E) Number of BM HSPC precursor in recipients of young FA donor.
(F) Ageing *Fancc* donor BM HSPC frequency analysis. N = 3 for Fancc^+/+^, N = 3 for Fancc^-/-^.
(G) Number of PB myeloid cell precursor in recipients of aging FA donor.
(H) Number of PB T cell precursor in recipients of aging FA donor.
(I) Donor chimerism in recipient BM of ageing *Fancc* donor. Each dot represents one animal. Each outlined dot represents the average value of a replicate containing multiple animals. Two-way ANOVA was performed. Recipient sample size = 7-10 mice per replicate, N =3 replicates per group.
(J) Number of BM HSPC precursor in recipients of aging FA donor.

For (C-E), (G-H) and (J), N =3 replicates per group; permutation test was performed; dashed lines connect the average precursor numbers. Error bars represent mean ± SD. nd, not determined; ns, non-significant; * p < 0.05, ** p < 0.001. # represents lowest p value possible for permutation test.

