## Supplementary figures and images for "Dynamic Tracking of Native Precursors in Adult Mice"

### Supplementary figures

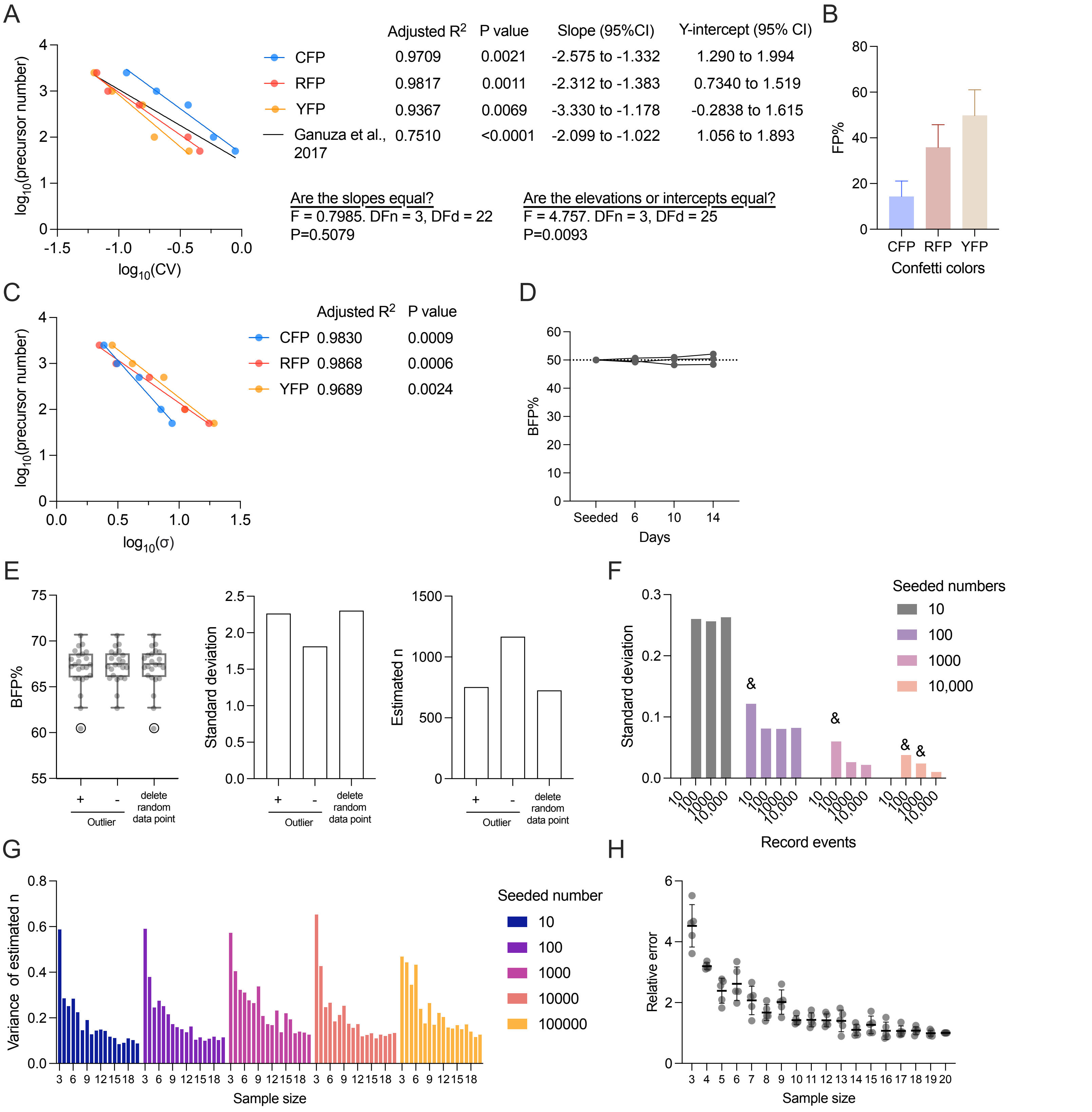

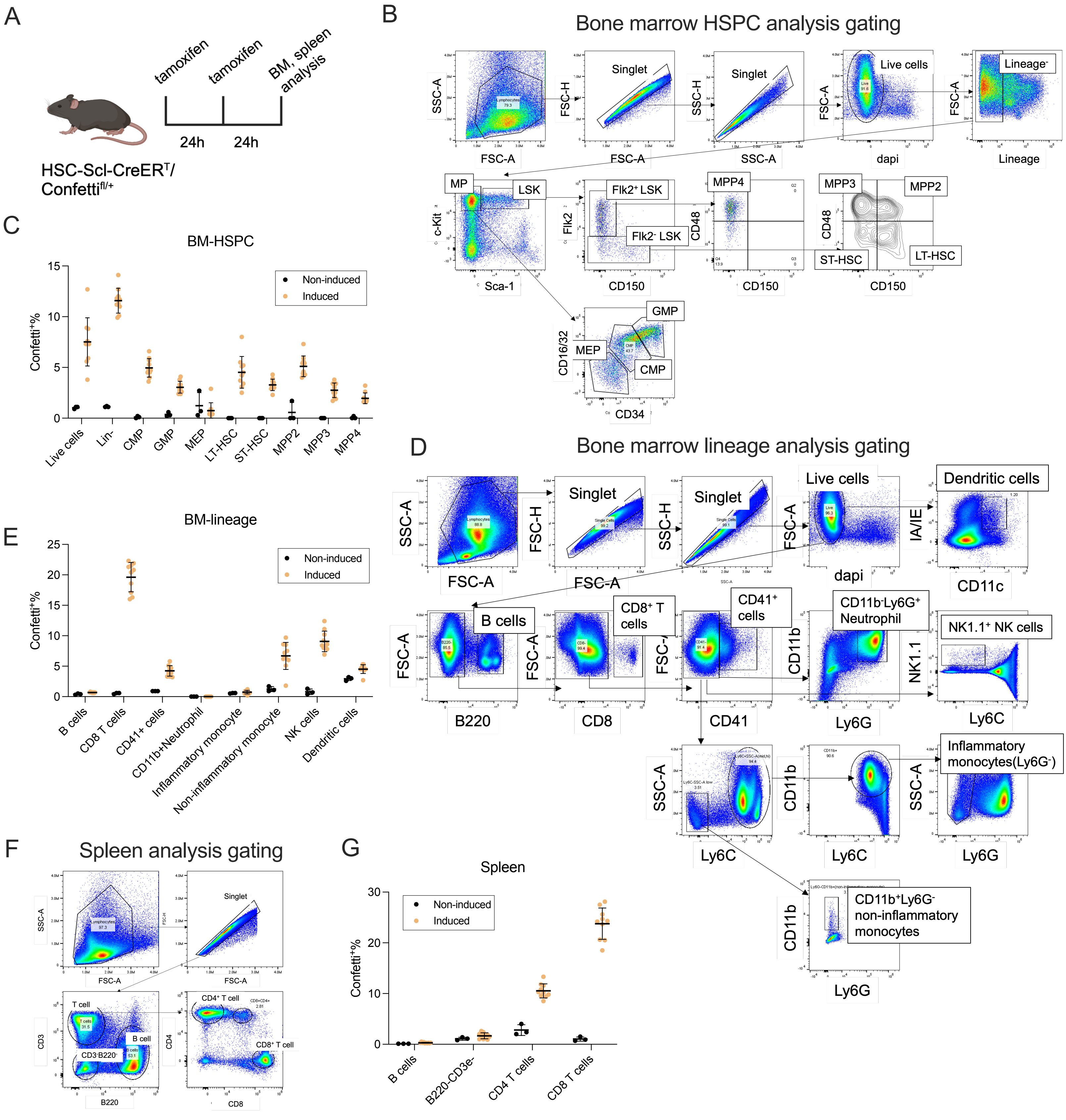

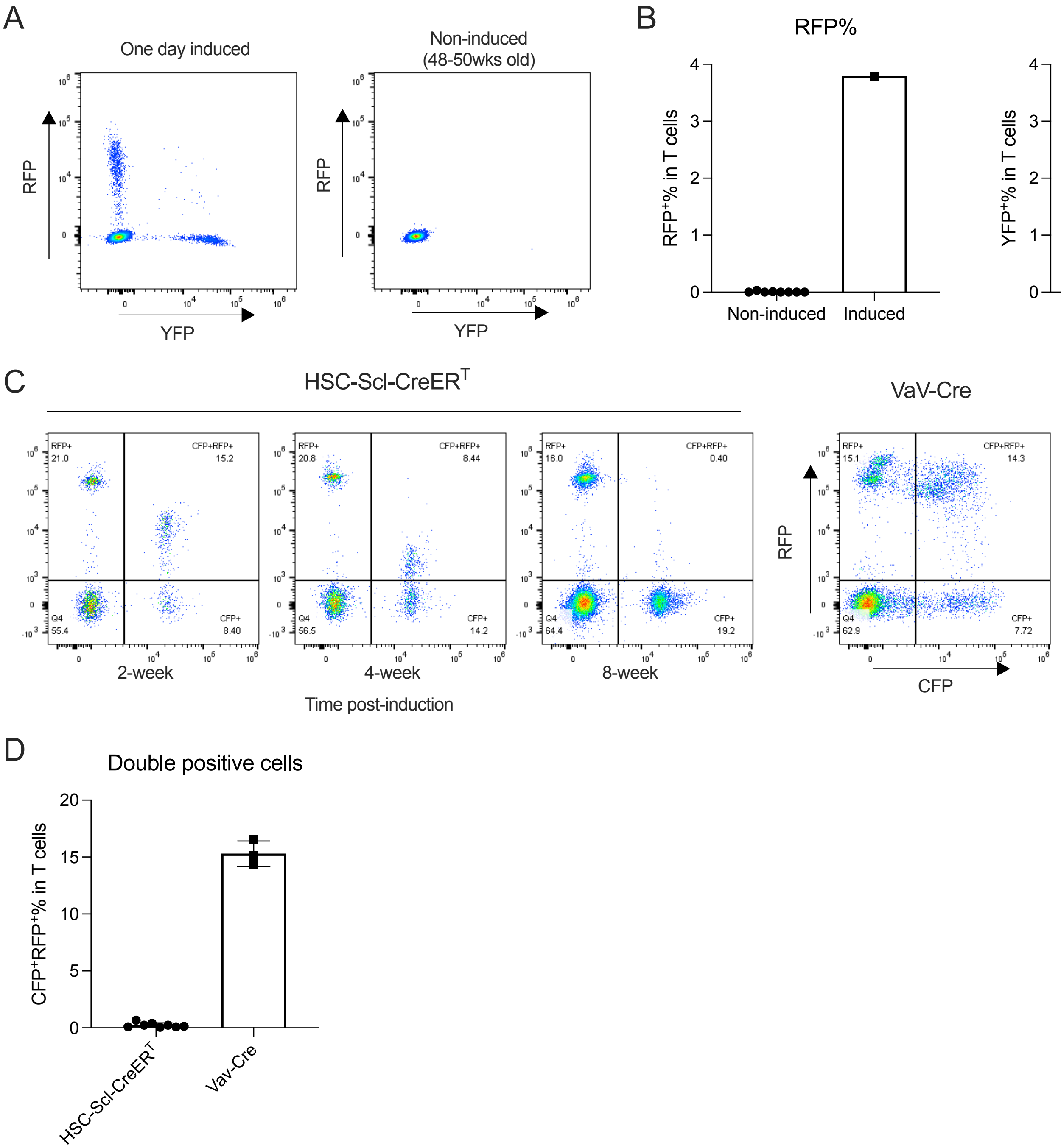

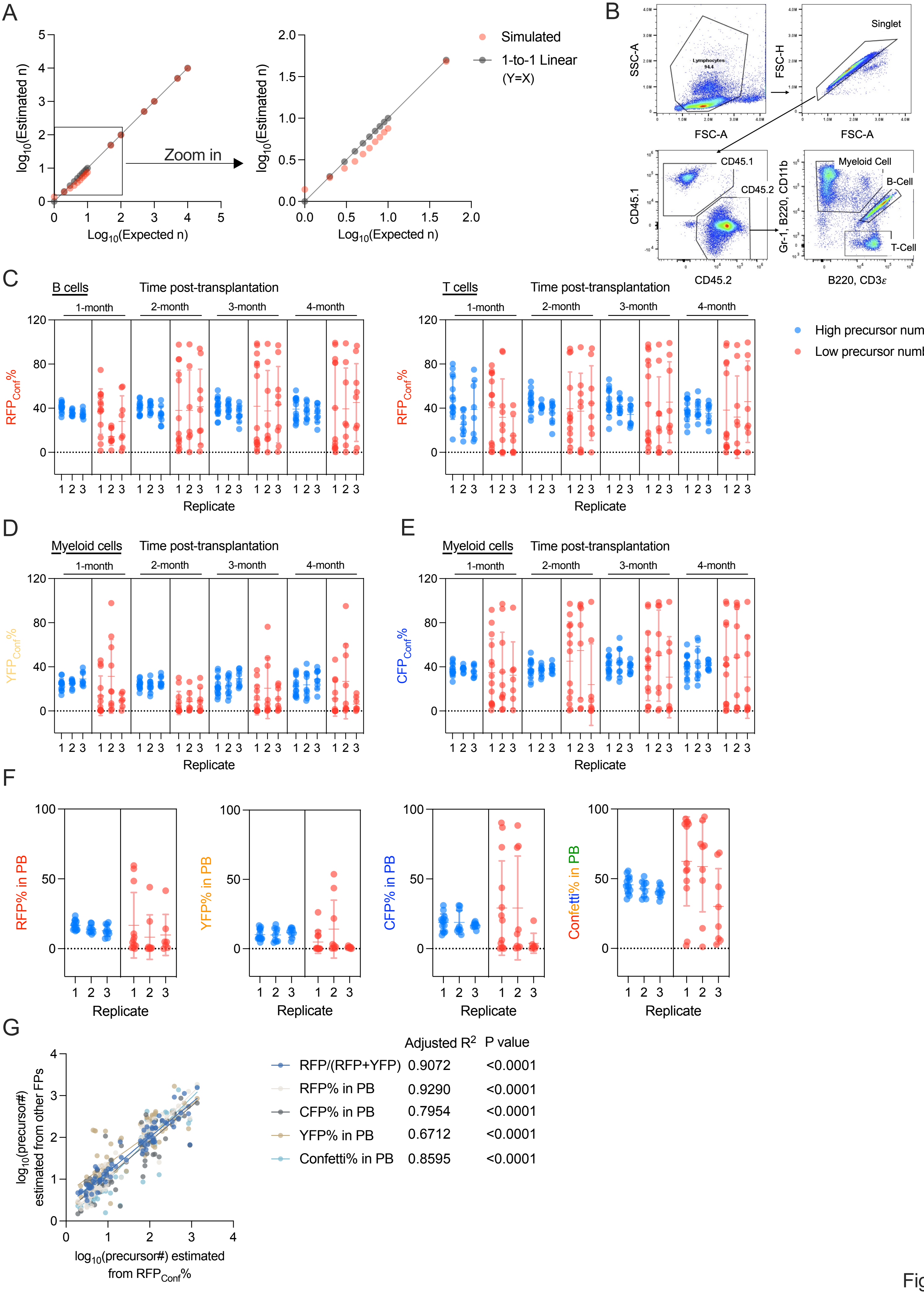

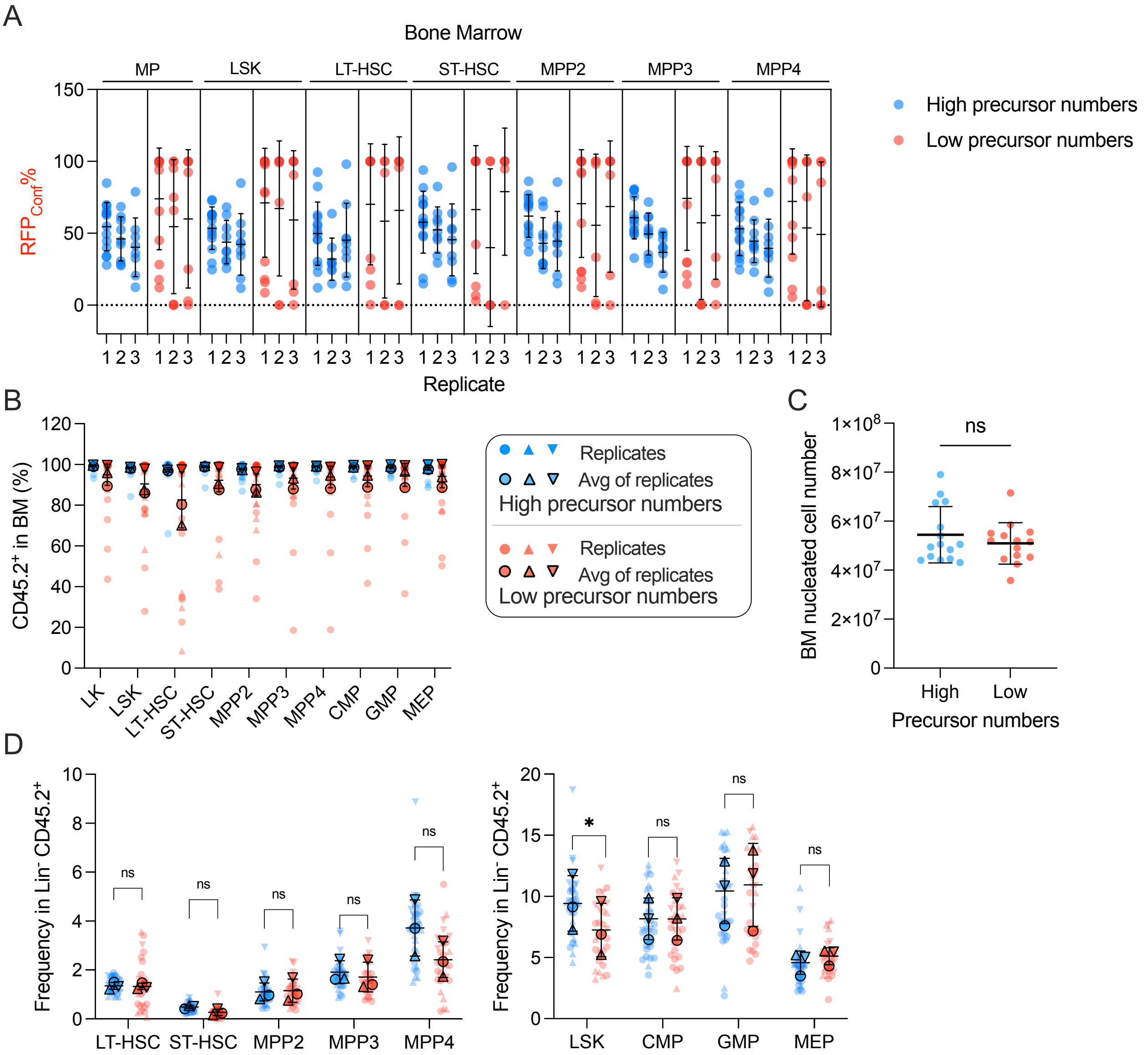

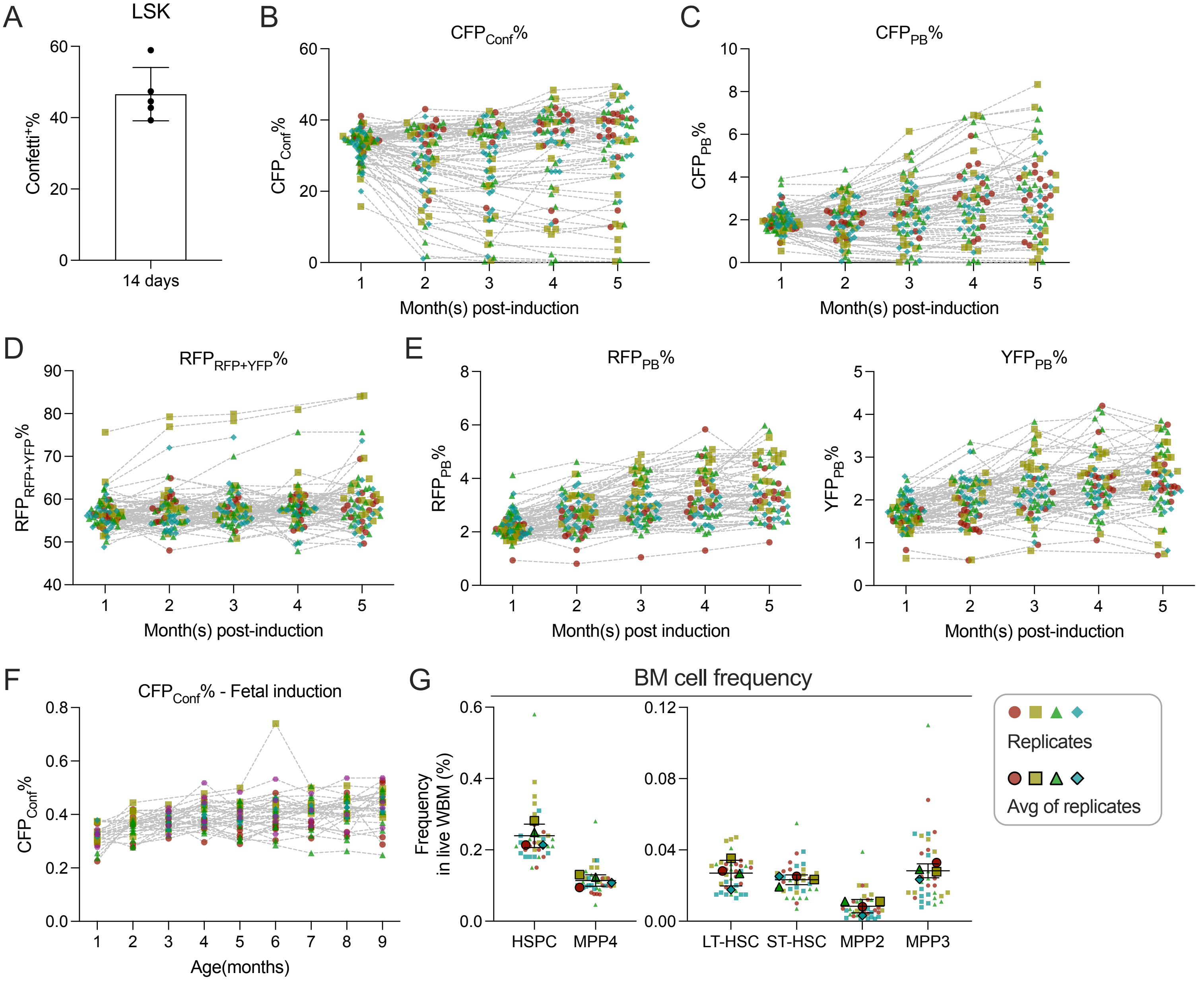

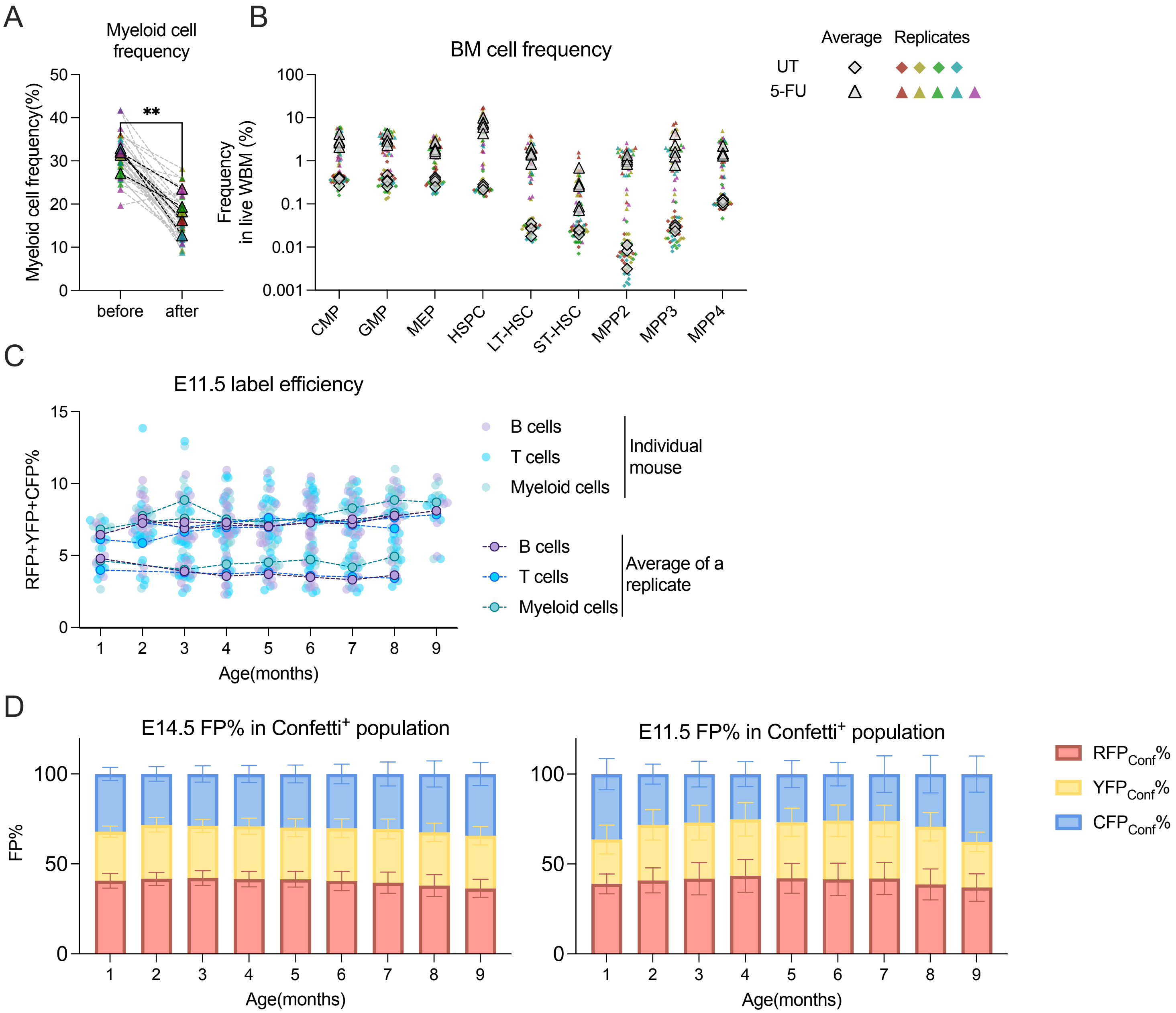

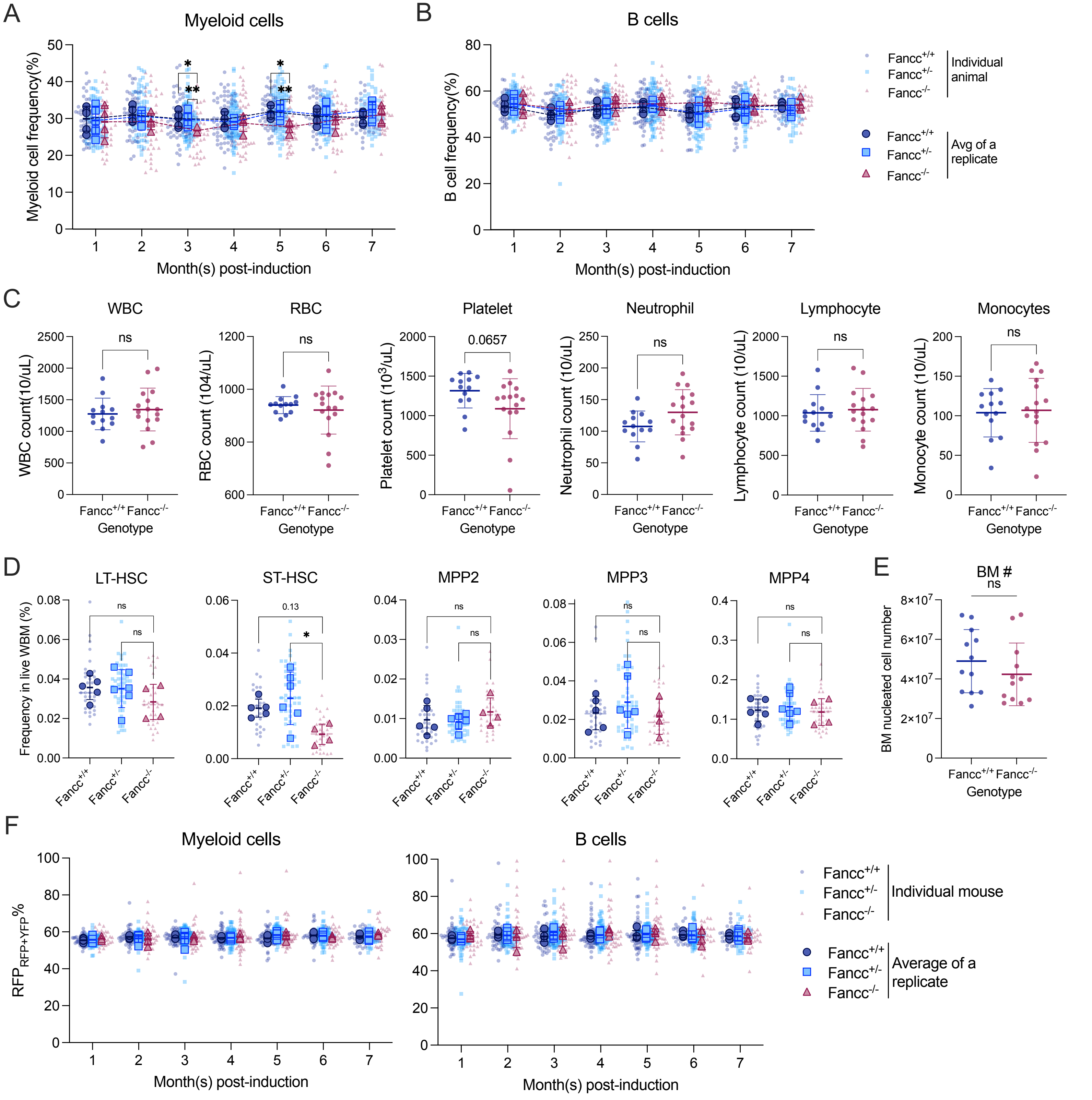

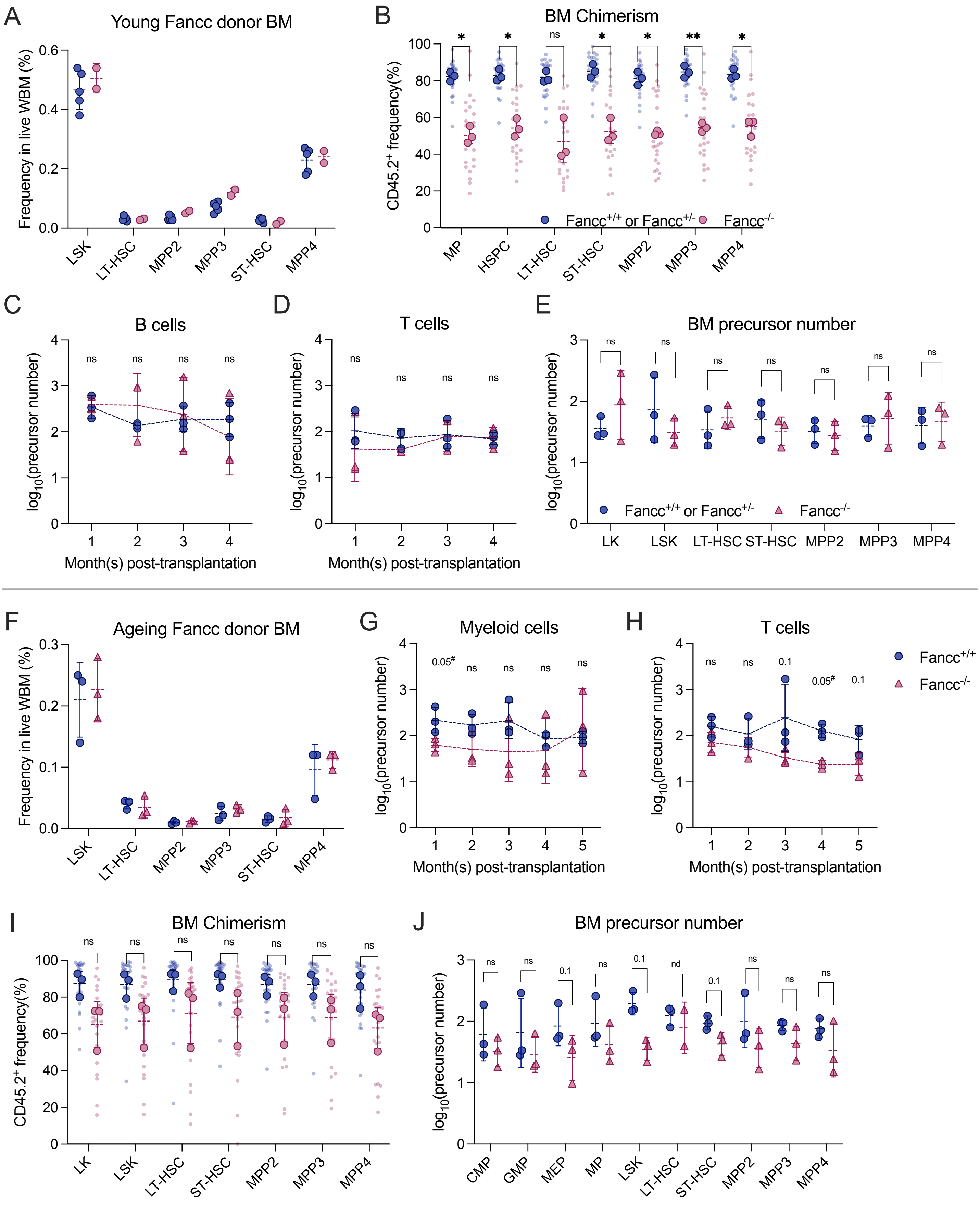
